# Understanding therapeutic tolerance through a mathematical model of drug-induced resistance

**DOI:** 10.1101/2024.09.04.611211

**Authors:** Jana L. Gevertz, James M. Greene, Samantha Prosperi, Natacha Comandante-Lou, Eduardo D. Sontag

**Affiliations:** Department of Mathematics and Statistics, The College of New Jersey, Ewing, NJ, United States; Department of Mathematics, Clarkson University, Potsdam, NY, United States; Department of Bioengineering, Northeastern University, Boston, MA, United States; Center for Translational & Computational Neuroimmunology, Columbia University Medical Center, New York, NY, United States; Department of Electrical and Computer Engineering, Northeastern University, Boston, MA, United States; Laboratory of Systems Pharmacology, Program in Therapeutic Science, Harvard Medical School, Boston, MA, United States

## Abstract

There is growing recognition that phenotypic plasticity enables cancer cells to adapt to various environmental conditions. An example of this adaptability is the persistence of an initially sensitive population of cancer cells in the presence of therapeutic agents. Understanding the implications of this drug-induced resistance is essential for predicting transient and long-term tumor tumor dynamics subject to treatment. This paper introduces a mathematical model of this phenomenon of drug-induced resistance which provides excellent fits to time-resolved *in vitro* experimental data. From observational data of total numbers of cells, the model unravels the relative proportions of sensitive and resistance subpopulations, and quantifies their dynamics as a function of drug dose. The predictions are then validated using data on drug doses which were not used when fitting parameters. The model is then used, in conjunction with optimal control techniques, in order to discover dosing strategies that might lead to better outcomes as quantified by lower total cell volume.

## 1 Introduction

The ability of cells and tissues to alter their response to chemical and radiological agents is a major impediment to the success of therapy. Such altered response to treatment occurs in infections by bacteria, viruses, fungi, and other pathogens, as well as in cancer [1–4]. Despite advances in recent decades, this reduction in the effectiveness of treatments, broadly termed *drug resistance*, remains poorly understood, and in some circumstances is thought to be inevitable [5].

One of the most clinically important examples of drug resistance is that involving the treatment of cancer via chemotherapy and targeted therapies. Drug resistance is a primary cause of treatment failure, with a variety of molecular and microenvironmental causes already identified [6]. For example, the upregulation of drug efflux transporters, enhanced DNA repair mechanisms, modification of drug targets, stem cells, irregular tumor vasculature, environmental pH, immune cell infiltration and activation, and hypoxia have all been identified as mechanisms which may inhibit treatment efficacy [4, 6, 6–14]. A vast amount of experimental and mathematical research continues to shed light on our understanding drug resistance. An excellent “roadmap” article (which also includes references to a number of relevant mathematical models) summarizes state-of-the-art approaches to understanding the role of both non-genetic plasticity and genetic mutations in cancer evolution and treatment response [15], while also highlighting current and future challenges.

Aside from understanding the mechanisms by which resistance to therapies may manifest, a fundamental question is *when* resistance arises. With respect to the initiation of therapy, resistance can be classified as either *pre-existing* or *acquired* [6]. The term pre-existing (also known as *intrinsic*) drug resistance is reserved for the case when the organism contains a sub-population (or a tumor contains a sub-clone) which resists treatment prior to the application of the external agent. Examples of the presence of extant resistance inhibiting treatments are abundant in bacteria and cancer, including genes originating from phyla Bacteroidetes and Firmicutes in the human gut biome [16], BCR-ABL kinase domain mutations in chronic myeloid leukemia [17, 18], and MEK1 mutations in melanoma [19]. Conversely, acquired resistance describes the phenomenon in which resistance first arises during the course of therapy from an initially drug-sensitive population. The question of whether resistance is pre-existing or acquired is a classical one in the context of bacterial resistance to a phage [20].

The study of acquired resistance is complicated by the question of *how* resistance emerges. Resistance can be *spontaneously* (also called *randomly*) acquired during treatment as a result of random genetic mutations or stochastic non-genetic phenotype switching. These cells can then be selected for in a classic Darwinian fashion [21]. Resistance can also be *induced* (or *caused*) by the presence of the drug [21–27]. These cells are often referred to as “drug tolerant persisters” in the literature [28]. That is, the drug itself may promote, in a “Lamarckian sense”, the (sometimes reversible) formation of resistant cancer cells so that treatment has contradictory effects: it eliminates cells while simultaneously upregulating the resistant phenotype, often from the same initially-sensitive (*wild-type*) cells.

The fundamentally distinct scenarios in which resistance may arise are illustrated in Fig. 1. Note that in each of the three cases (pre-existing, spontaneous acquired, and induced acquired), the post-treatment results are identical: the resistant phenotype dominates. However, the manner by which resistance is generated in each case is fundamentally different, and the transient dynamics (both drug-independent and drug-dependent) may vary drastically in each scenario. As a result, the same therapeutic design may result in different outcomes depending on the (dominant) mode of resistance. This “phenotypic plasticity” allows cancer cells to adapt to different environmental circumstances [29]. Although there is experimental evidence for these three forms of drug resistance, differentiating them experimentally is non-trivial. For example, what appears to be drug-induced acquired resistance may simply be the rapid selection of a very small number of pre-existing resistant cells, or the selection of cells that spontaneously acquired resistance [21, 30]. It is in this realm that mathematics can provide invaluable assistance. Formulating and analyzing precise mathematical models describing the previously mentioned origins of drug resistance can lead to novel conclusions that may be difficult, or even impossible, given current technology, to determine utilizing experimental methods alone. The main goal of this work is to validate a previously developed mathematical model of drug-induced resistance, and to explore the implications for protocol design.

**Figure 1:**
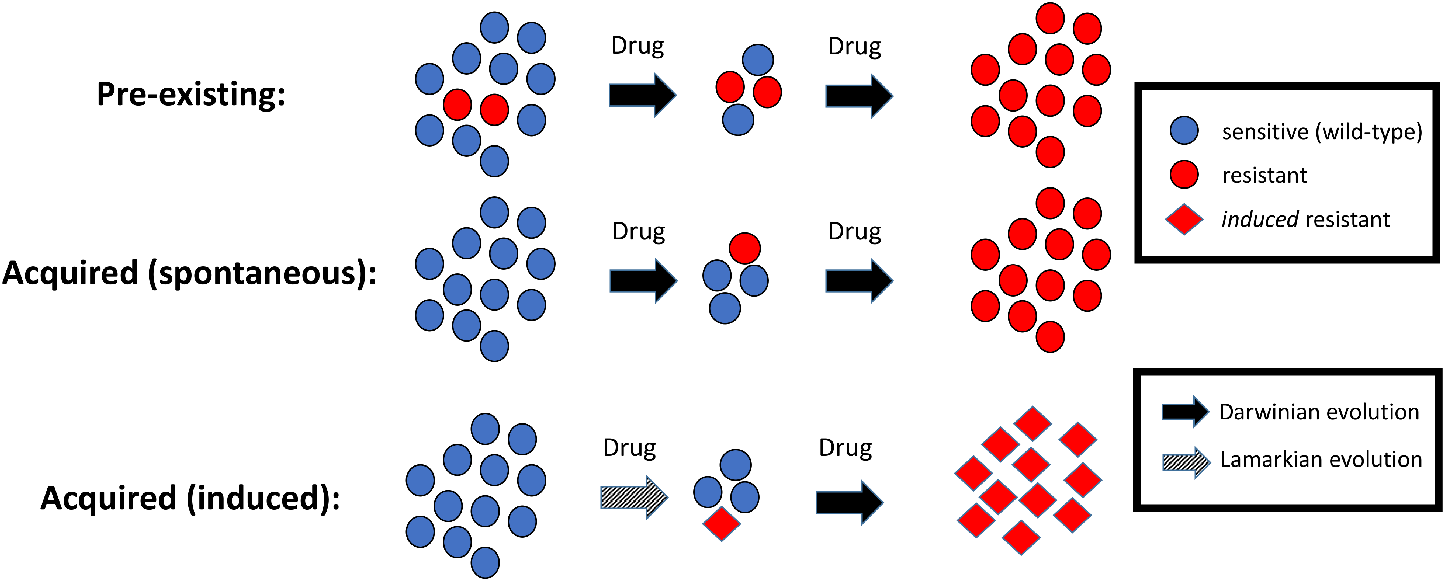
Three distinct ways in which drug resistance may be generated. In the first row, resistance is preexisting, so that resistant clones are selected during therapy. In spontaneously acquired resistance, random genetic and epigenetic modifications produce a resistant phenotype during therapy, where again selection dominates. In the case of induced acquired resistance, the application of the drug *causes* the resistant cell to arise, so that the mechanism is *not* simply random variation (e.g. mutations) followed by classical Darwinian selection. Instead, there is a “Lamarckian” component (grey arrow). Note that any combination of these three mechanisms may be causing resistance in a single patient.

In Section 2 we describe a set of experimental data with strong evidence of resistance induction [31, 32], which will form the basis of our work. In Section 3 we will present a preliminary mathematical model based on our original resistance model from [33], and discuss biologically motivated changes, followed by a discussion of the methodology that we utilize to fit the data and analyze the goodness-of-fit. In Section 4 we demonstrate that our modified model gives excellent fits across a range of drug doses, show that a more detailed model does not improve fits, and perform a cross-validation study to confirm the model by testing it on doses not used in fitting parameters. We also study optimal dosing in this section. Finally, Section 6 has concluding remarks and directions for future work.

## 2 Motivating Experimental Data

Melanomas harboring mutations in BRAF, which result in excessive growth signaling, are commonly treated with small-molecule BRAF inhibitors such as vemurafenib, dabrafenib, and encorafenib [34]. These drugs prevent the over-activation of pathways controlled by BRAF, including several types of MAPK pathways. As with other therapeutics, resistance to BRAF inhibitors poses a significant barrier to treatment success. Drug-induced reprogramming by vemurafenib has been well-studied experimentally [35].

In an effort to better understand resistance to vemurafenib, Fallahi-Sichani et al. subjected COLO858 melanoma cells (which harbor a BRAFV600E mutation) *in vitro* to vemurafenib at various drug concentrations before imaging the cells over the course of 84 hours [31]. Single cell imaging during vemurafenib exposure revealed three possible outcomes: death, quiescence (cells survive but do not divide), and resistance (cells survive and divide). Interestingly, if the COLO858 cells were pre-treated overnight with a low-dose of vemurafenib before their entrance into the higher-dosed assay conditions, greatly increased cell viability was observed [31]. This finding indicates that a lower dose of drug may “teach” tumor cells the mechanism of resistance, which will then allow them to grow under treated conditions that they would not have otherwise survived. They also found that this vemurafenib resistance is reversible when non-sensitive cells were plated in fresh drug media. Upon the return of vemurafenib to these sorted cells’ culture conditions, resistance was re-induced, indicating that this mechanism of resistance is both reversible and persistent [31].

In follow-up work, COLO858 cells were studied *in vitro* under varying vemurafenib concentrations over a period of 120 hours [32]. An algorithm was used to count the number of cells, division and death events over this time period. As in Fallahi-Sichani et al. [31], cells were observed to be in either a sensitive, quiescent, or resistant state. Using this cell count data, the authors calculated the instantaneous death and growth rate of COLO858 cells while subjected to treatment with various doses of vemurafenib [32]. We use the 0.032uM, 0.1uM, 0.32uM, 1uM, and 3.2uM datasets from [32] in our paper.

## 3 Model and Computational Methods

In an effort to better understand the role of selection and induction in the evolution to drug resistance, in [33] we introduced a simple, phenomenological mathematical framework to distinguish between spontaneous and induced resistance. Specifically, we initially considered a tumor population consisting of sensitive (*S*), non-reversible resistant (*R*_*n*_), and reversible resistant (*R*_*r*_) cells, whose dynamics are described by the following set of ordinary differential equations (ODEs):

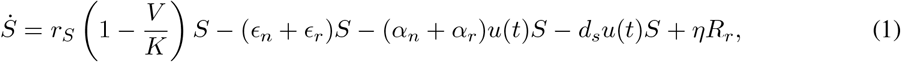

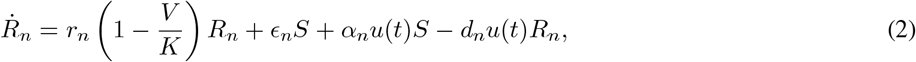

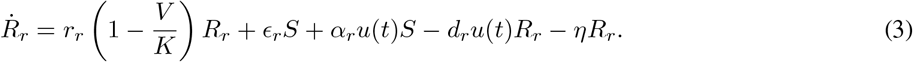

Here *u*(*t*) represents the effective applied drug dose at time *t*, and is thought of as a control input, and *V* is the total tumor population (*V* := *S* + *R*_*N*_ + *R*_*r*_). Dots represent time derivatives. By definition, the drug is less effective against resistant cells, so we assume that *d*_*n*_, *d*_*r*_ ≤ *d*. We also assume that *r*≥*r*_*n*_, *r*_*r*_, as experimental evidence supports that resistant cells grow slower than nonresistant cells [36, 37]. All cells compete for resources via a joint carrying capacity *K*, and resistance may develop in *both* a spontaneous drug-independent (*ϵ*_*n*_*S, ϵ*_*r*_*S*) and an induced drug-dependent (*α*_*n*_*u*(*t*)*S, α*_*r*_*u*(*t*)*S*) manner. All rate parameters in the above system are assumed nonnegative.

In equations (1) - (3), non-reversible cells *R*_*n*_ can be thought of as resistant cells that arise via genetic mutations. As “undoing” genetic mutations are unlikely, reversibility is assumed negligible. The reversible cells *R*_*r*_, on the other hand, are resistant cells that arise via phenotype switching. As this process is reversible [21, 24], we include a re-sensitization term (*ηR*_*r*_) for this subpopulation. The drug induction terms *α*_*n*_*u*(*t*)*S* and *α*_*r*_*u*(*t*)*S* assume the log-kill hypothesis, in which the rate of induced resistance is proportional to the applied dosage [38].

We have previously studied the effects of resistance induction in a simplified version of this model, where we assumed that the reversible and non-reversible resistant cells exhibit identical kinetics. Specifically, we consider the possibility that *ϵ*_*n*_ = *α*_*n*_ = 0 in (3) and thus simplified the resistant dynamics to a single population, assuming that there were no non-reversible resistant cells in the population initially (*R*_*n*_(0) = 0). Thus, with *R* := *R*_*r*_ and *ϵ* := *ϵ*_*r*_, *α* := *α*_*r*_, *r*_*R*_ := *r*_*r*_, and *d*_*R*_ := *d*_*r*_, we proposed the below two-dimensional system:

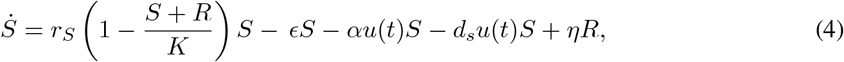

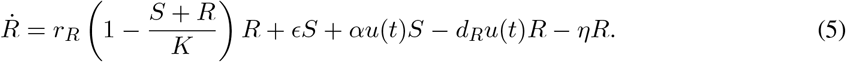

Using this simplified model, we found a *qualitative* distinction in tumor response to two standard treatment schedules (continuous versus pulsed) when the drug has the same cytotoxic potential but different levels of induction (i.e. identical *d*_*S*_ and *d*_*R*_, but different *α* in equations (4) - (5)). In particular, for a fixed set of model parameters, when a drug cannot induce resistance (*α* = 0), constant therapy significantly outperforms pulsed therapy, with the time to treatment failure being nearly seven times longer using constant therapy. However, when a drug can induce resistance (*α >* 0), pulsed therapy improves the duration of therapeutic efficacy when compared to constant treatment [33]. This illustrates how the mechanism of drug resistance can strongly influence response to a treatment protocol. A rigorous analysis of the optimal control structure (both theoretical and numerical) can be also be found in [39].

As discussed below, the originally proposed system (4) (5) cannot qualitatively capture key features of the experimental data in [32]. Thus, in Section 3.1, we propose a modification to this original model, which includes time delays with respect to the effective dose. We also discuss the precise methodology utilized for model fitting and validation in Section 3.2, and we conclude this section with a description of the numerical optimal control formulation for time-varying doses in Section 3.3. All code is available at https://github.com/sontaglab/Induced_Resistance.

### 3.1 Revised Mathematical Model

The initial increase in the number of cells at all doses of vemurafenib considered in [32] (see Figure 2(a), for example) indicates that there is a delay in drug action. Indeed, the response the control and to all five administered vemurafenib doses (0.032, 0.1, 0.32, 1, 3.2 *µ*M) are indistinguishable in Figure 3(a) in [32] for approximately the first 24 hours of the experiment. Biologically, this indicates that there is a delay in drug effect, which may be due to various factors such as drug uptake, engagement of the target, initiation of the apoptotic pathway, or cell-cycle inhibition. Note that model (4) - (5) does not include any such delay, and thus any applied difference in dosage will result in an immediate response by the cell population.

**Figure 2:**
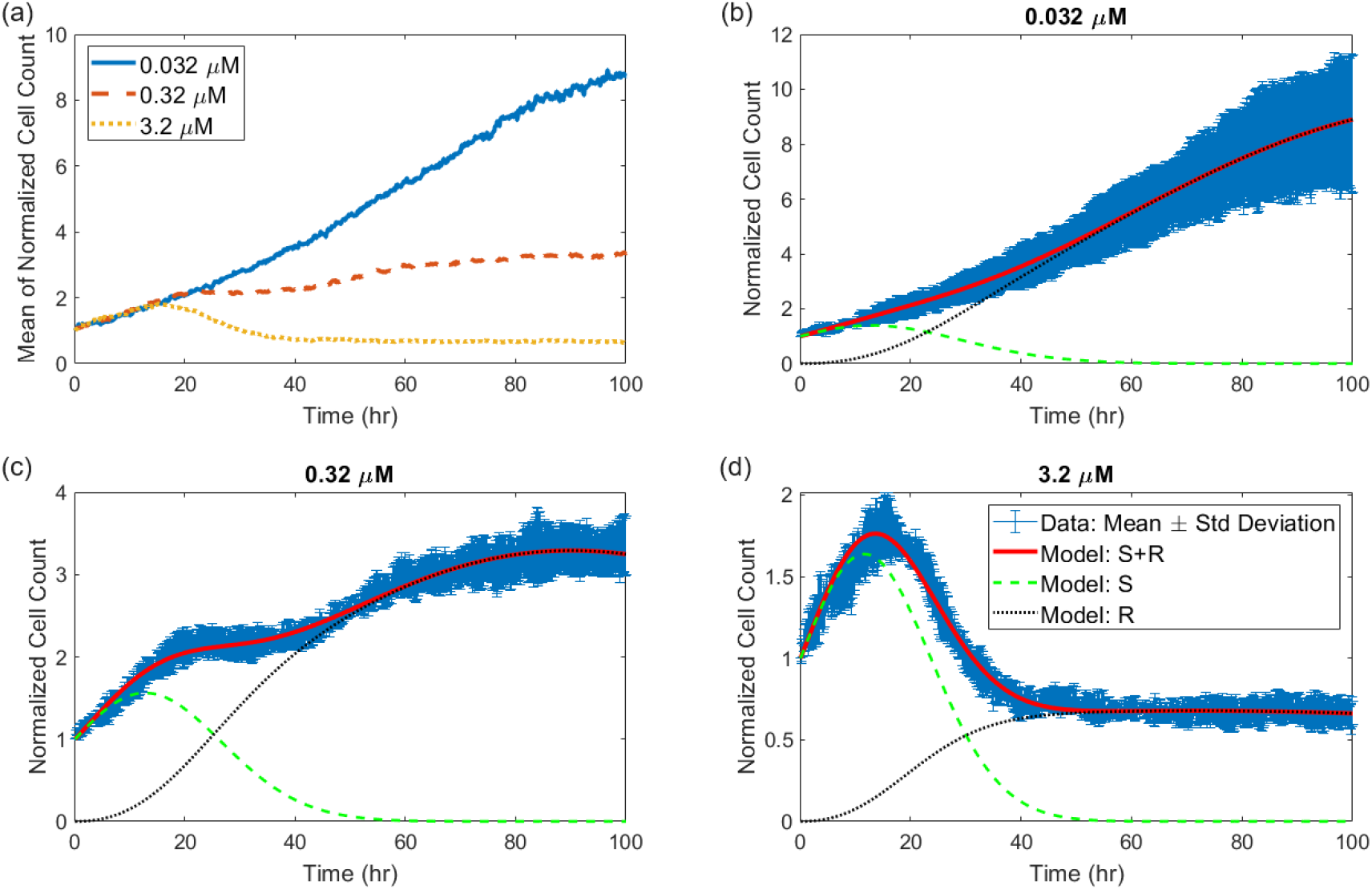
Data and fits for the two-population model. (a) Mean across four replicates of the normalized cell count data from [32] used for model fitting. Best model fits are shown at a drug concentration of (b) 0.032 *µ*M, (c) 0.32 *µ*M, and (d) 3.2 *µ*M. Standard deviations (four replicates) are shown in blue in (b)-(d). The fitting process results in time course predictions for the respective sizes of the sensitive (*S*) and resistant (*R*) cell subpopulations.

**Figure 3:**
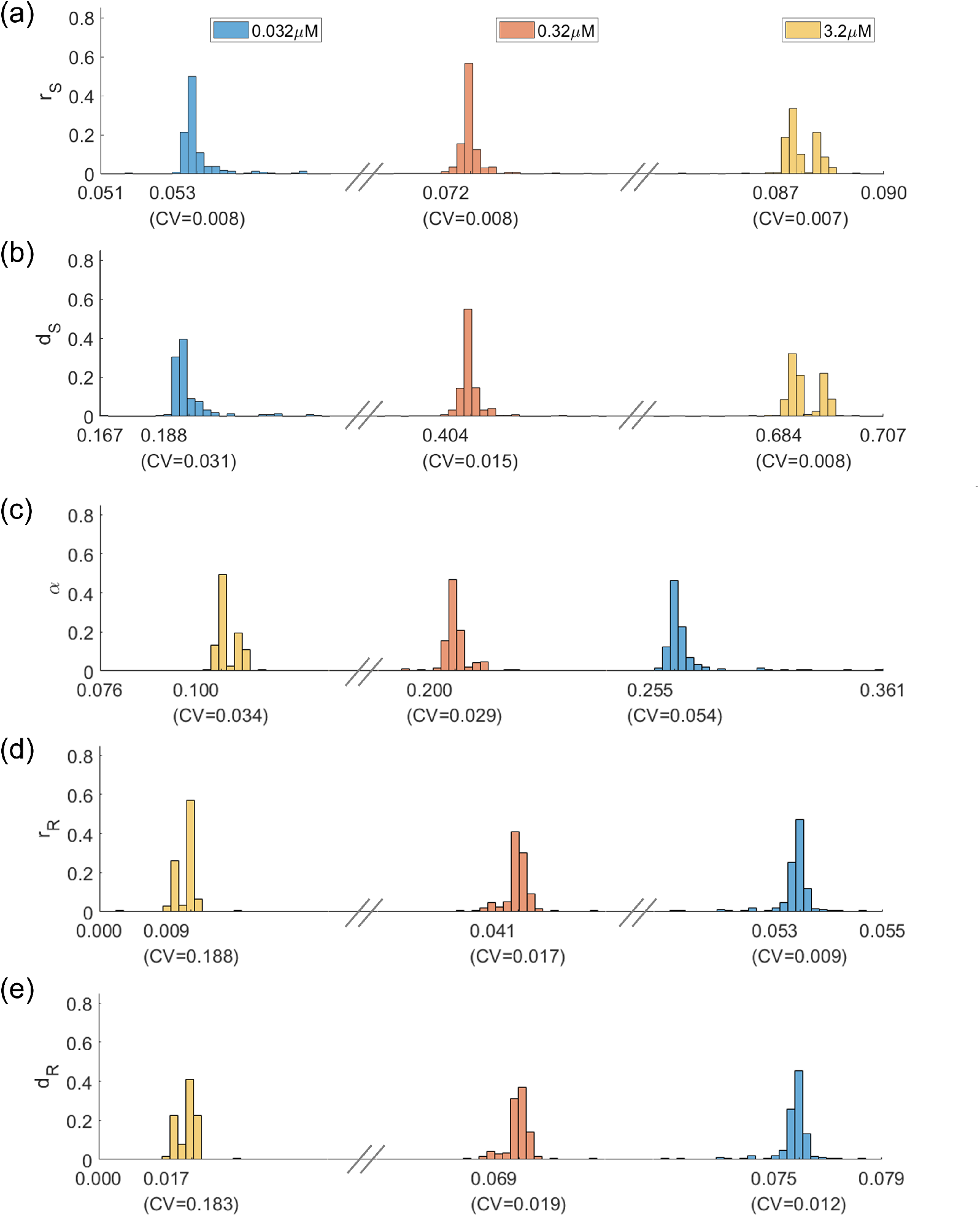
Practical identifiability of parameters in two-population model. Distribution of parameters that yield model fits within 5% of the optimal parametrization for three doses: 0.032 *µ*M in blue, 0.32 *µ*M in red, and 3.2 *µ*M in yellow. Below each distribution, the mean is provided, along with the coefficient of variation (CV). Plots are shown for (a) *r*_*S*_, (b) *d*_*S*_, (c) *α*, (d) *r*_*R*_, and (e) *d*_*R*_.

For this reason, we propose a modified version of our model that incorporates a delay on both drug-induced cell kill and drug-induced resistance. Specifically, we differentiate between the applied dosage *u* and the effective dosage *v*; here *u* represents the drug concentration of vemurafenib in the *in vitro* experiment, while *v* corresponds to the phenotypic effect of the dosage on cell kill and induction rates. We assume that the dynamics on the rate parameters can be described via classical input (*u*) tracking [40], with constant rates *γ*_1_ and *γ*_2_ determining the delay timescales for the drug-induced cell death and resistance-induction rates, respectively. The model we consider takes the following form:

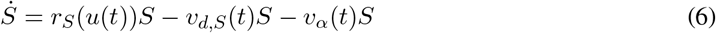

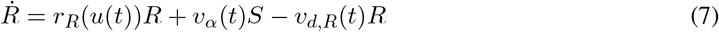

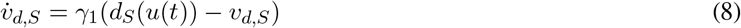

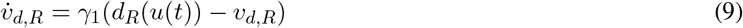

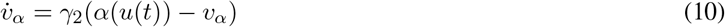

As in (4) - (5), state *S* represents the (normalized) number of sensitive cells and *R* represents the normalized number of cells that have acquired an “inducible drug resistant” state. The parameters *r*_*S*_ and *r*_*R*_ represent growth rates of the drug sensitive and drug resistant populations, respectively, as a function of the applied dose *u*(*t*). As opposed to parameters, we now consider *states v*_*d,S*_ and *v*_*d,R*_, which represent each subpopulations’ drug-induced death rate, and *v*_*α*_, the rate at which drug sensitive cells are induced to transition to the resistant phenotype. Note that division rates *r*_*S*_ and *r*_*R*_ are assumed to respond to *instantaneous* changes in the applied dosage *u*, while the *effective* apoptosis and induction rates *v*_*d,S*_, *v*_*d,R*_, and *v*_*α*_ approach their asymptotic values at a rate determined by *γ*_1_ and *γ*_2_, respectively. This tracking dynamic thus models the observed delay in the onset of observable effects of the applied chemotherapy; see Section 3.2 for further discussion of the values utilized for the delay timescales *γ*_1_ and *γ*_2_. We emphasize that all rate parameters (*r*_*S*_, *r*_*R*_, *d*_*S*_, *d*_*R*_ and *α*) are dependent on the applied dose *u*(*t*), and we denote this via expressions of the form *r*_*S*_(*u*(*t*)), *r*_*R*_(*u*(*t*)), etc. As to be discussed in Section 3.2, these rates will be estimated from the normalized cell count data for different constant doses *u*(*t*) ≡ *u*_constant_.

Although a carrying capacity was assumed in our original model (4) - (5), here we consider the simpler scenario of exponential growth. This is because our original model was envisioning an *in vivo* setting, whereas herein we are modeling *in vitro* data under non-limiting growth conditions. In this model, we also assume that spontaneous (genetic) resistance is negligible on the short time scales considered. This is equivalent to setting *ϵ* = 0 in the original model. Given that the data the model is being fit to counts of cell number *only in the presence of drug*, we also assume that re-sensitization of resistant cells is negligible, and thus set *η* = 0 in equations (4) - (5).

As an example of the dynamic introduced by rate tracking in cell death and induction, consider equation (8) describing the effective apoptosis rate for the sensitive population:

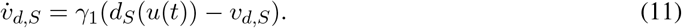

Assuming *v*_*d,S*_(0) = *v*_0_, the solution of this linear ODE is given by the variation of parameters formula

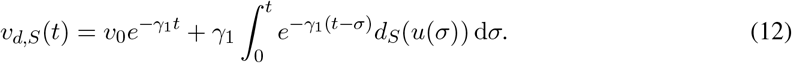

Similar statements hold for *v*_*d,R*_ and *v*_*α*_. Hence, equations (6) - (10) represent a time-varying two-dimensional system in the *S* and *R* populations.

The experimental data from [32] utilized in this work considers *constant* applied doses *u*, so that when fitting model (6) - (10) to experimental data, we assume that *u*(*t*) ≡ *u*, a constant dose. Thus, we suppress the explicit applied dosage dependence of the rate parameters, and write *r*_*S*_ := *r*_*S*_(*u*), *r*_*R*_(*u*) := *r*_*R*_(*u*), *d*_*S*_ := *d*_*S*_(*u*), *d*_*R*_ := *d*_*R*_(*u*), and *α* := *α*(*u*), as these are constants (but are dose-dependent). The expressions in (12) thus simplify as follows. Assuming that the effective dosages for the cell kill and induction rates are zero initially (i.e. *v*_*d,S*_(0) = *v*_*d,R*_(0) = *v*_*α*_(0) = 0), we see from (12) that the solutions of equations (8) - (10) take the form

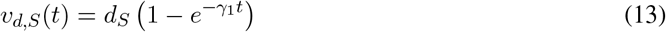

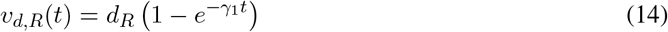

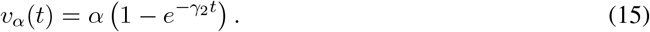

Using the above, system (6) - (10) becomes

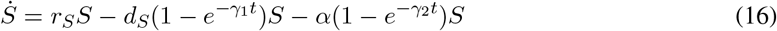

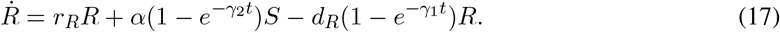

The above system is what is used to determine the rate parameters as a function of (constant) dose *u*. Note that we will later consider time-varying doses *u* = *u*(*t*) when formulating and analyzing an optimal control problem (see Sections 3.3 and 4.4); in this case, we will consider the original modified model (6) - (10).

### 3.2 Fitting Methodology

The *in vitro* data from [32] includes five different dosing regimes covering a wide range of drug concentrations. Three of those doses were selected to parameterize the model in eqns. (16)-(17): 0.032 *µ*M, 0.32 *µ*M, and 3.2 *µ*M. The doses of 0.1 *µ*M and 1 *µ* are withheld for validation purposes. All model parameters are fit to this data *except* the delay terms *γ*_1_ and *γ*_2_. Data in [41] suggests that the onset of drug-adapted conditions for the A375 BRAF V600E melanoma cell line is observed at around 24 hours. In another study [42], cytotoxic effects reached full effect between 120 and 130 hours. Although these studies were not specifically related to COLO858 cells, and the exact times are approximations, to decrease the complexity of the model (and specifically parameter estimation), we decided to fix *γ*_1_ = *γ*_2_ = 0.01. This corresponds to both terms achieving roughly 50% of their maximal effect at 72 hours. Given that we assume that all resistance is drug-induced, and that *S* and *R* represent *normalized* cell counts, the model is always solved using the initial conditions of *S*(0) = 1 and *R*(0) = 0.

The above assumptions result in needing to fit five model parameters per drug dose: the sensitive cell growth rate *r*_*S*_, the sensitive cell drug-induced death rate *d*_*S*_, the drug-induced resistance rate *α*, the resistant cell growth rate *r*_*R*_ (assumed to be ≤ *r*_*S*_), and the resistant cell drug-induced death rate *d*_*R*_ (assumed to be ≤ *d*_*S*_). We seek to minimize the sum of the absolute difference between the model and the data:

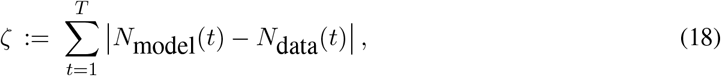

where *N*model(*t*) = *S*(*t*) + *R*(*t*) is the model-predicted normalized cell count at time *t*, and *N*data(*t*) is the normalized experimental mean cell count at time *t* (see Fig 2(a)).

To parameterize the model for each drug dose, a multi-start algorithm is implemented. Multi-start fitting methods repeat the optimization process many times from different starting points, seeking to mitigate the risk of getting stuck at local minima and improve the chances of finding the globally optimal parameter set. The optimization algorithm utilized for each starting point is MATLAB’s *fmincon* function, which executes an interior-point method for solving constrained minimization problems. Beyond the inequality constraints posed on growth and drug-induced death rate of resistant cells, a restriction was set on the maximum number of function evaluations (see Table A.1). Further, positive (though very small) lower bounds were imposed on all parameters to avoid numerical integration issues in solving the model in equations (16)-(17). Upper bounds, also detailed in Table A.1, were imposed to ensure that each parameters remained within a biologically-plausible range.

The constrained optimization problem was solving using *fmincon* for *K* = 1000 starting parameter guesses. Each starting parameter guess is obtained by using a quasi-Monte Carlo method to randomly sample the parameter space. The lower and upper bound for each parameter was set to the constraints imposed on the optimization problem itself. We particularly used Sobol’s low-discrepancy sequences, which have uniformity properties other samples techniques lack while being computationally efficient [43]. The multi-start fitting algorithm thus starts by randomly sampling *K* Sobol points of the form (*p*_1_, …, *p*_*n*_), where *n* is the number of parameters. As each *p*_*i*_ in a sampled point are in the range [0, 1], we then scale each values of *p*_*i*_ to be in the range defined by its lower and upper bound. For each such parameterization, *fmincon* is implemented using that parameter set as the starting guess.

The end result of this multi-start algorithm is a set of *K* “best-fit” parameterizations of the model. The parameterization with the lowest cost function value *ζ*_*opt*_ is selected as the optimal model parameterizaton. Any other “best-fit” parametrization for which *ζ* varies from *ζ*_*opt*_ by 5% or less is considered a “suboptimal” parameter set. We will use these suboptimal parameter sets to study the sensitivity and identifiability of the model parameters.

### 3.3 Optimal control formulation

Although the experimental data and model was calibrated to constant doses, we are interested in utilizing our mathematical model to investigate dosing strategies which minimize the cancer cell population (*S* + *R*) at the end of the experiment; see Section 4.4 for further motivation. That is, we numerically compute optimal time-varying applied dosing strategies *u*(*t*) which minimize *S*(*t*_*f*_) + *R*(*t*_*f*_), where *t*_*f*_ is a fixed final time. The precise formulation of this optimal control problem is provided in this section.

By fitting model (16) - (17) to the experimental data, we obtain expressions for the parameters *θ*(*u*) := (*r*_*S*_(*u*), *d*_*S*_(*u*), *α*(*u*), *r*_*R*_(*u*), *d*_*R*_(*u*)) as a function of three doses (*u* = 0.032 *µ*M, 0.32 *µ*M, 3.2 *µ*M); as discussed in Section 4.3, these expressions were further validated on two intermediate doses (*u* = 0.1 *µ*M, 1 *µ*M). To obtain an expression for *θ* as a function of *all* doses in the continuous range [0.032, 3.2] *µ*M, we smoothly approximate the piecewise linear approximations used in prediction, and thus obtain functions *θ* = *θ*(*u*). (see Section A.3 and Figure A.2 for details). The optimal control problem is formulated utilizing these expressions for *θ*(*u*).

Denote by *x* := (*S, R, v*_*d,S*_, *v*_*d,R*_, *v*_*α*_) ∈ ℝ^5^ the state of model (6) - (10). The system then takes the general form 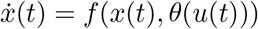, with

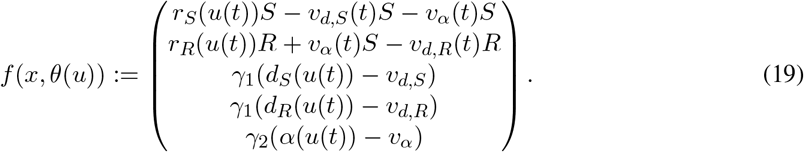

The optimal control problem is then defined on a finite-time horizon via the initial-value problem 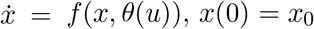 on [0, *t*_*f*_ ], with *t*_*f*_ fixed during each optimization. As in all simulations in this work (see Section 3.2), we fix the initial conditions as an entirely sensitive cell population and zero effective dose; the latter models drug delay factors as discussed in Section 3.1. Thus, *x*_0_ = (1, 0, 0, 0, 0).

We define our objective as the final cancer cell population size at *t*_*f*_,

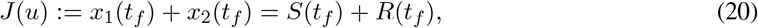

and we determine the dosing strategy *u*_*_ which minimizes *J*. Since our rate functions *θ*(*u*) are calibrated only for doses between 0.032 *µ*M and 3.2 *µ*M (see (30)), we restrict our control set *U* to the convex, compact interval *U* := [0.032, 3.2]. The set of admissible controls then takes the form 𝒰 := {*u* : [0, *t*_*f*_ ] → ℝ |*u* is Lebesgue measurable }. Recall that to guarantee the existence of a minimizing function *u*_*_ of *J*, we need to define the set of admissible controls as measurable (see, for example, [44]). However, we see below that this is not restrictive for the above problem considered in this work, and all optimal controls appear to be piecewise continuous (see, for example, Figure 7).

To incorporate toxicity constraints, we bound the total applied dosage by a constant *M* :

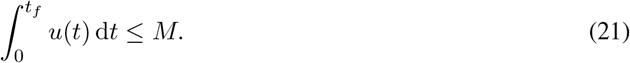

This constraint can be incorporated as an auxiliary state variable

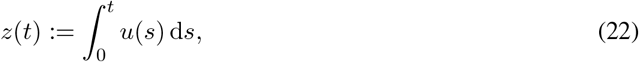

so that (21) becomes *z*(*t*_*f*_) ≤ *M*. The extended state of the original system defined by the vector field in (19) is thus defined as 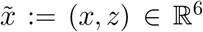, and the extended vector field is 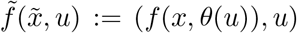. Since *z*(0) = 0, the initial conditions for 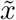 become 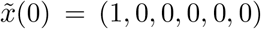. The control problem then incorporates an additional constraint on the last component of 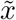 at the final time *t*_*f*_ :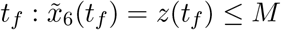.

In summary, the optimal control problem we analyze is attempting to minimize the final cancer cell population at final time *t*_*f*_,

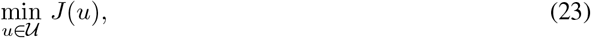

subject to the following initial-value problem in ℝ^6^

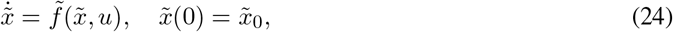

together with the additional final time constraint

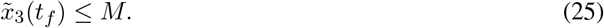

Lastly, we note that by construction, the rate functions *θ*(*u*) depend on the logarithm of the dose, i.e. *θ*(*u*) = *θ*(log_10_(*u*)) (see Section 4.3). Numerically, we thus reformulate the control with respect to the logarithm of the dose, so that we solve this system with respect to 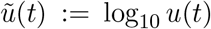. Effectively, this redefines the control set as 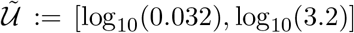, as well as the dynamics on the auxiliary variable *z* as 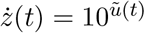. Thus, the vector field numerically integrated takes the form 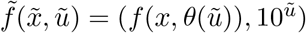.

## 4 Results and Discussion

### 4.1 Two-population model was able to adequately capture the dynamics of a wide range of dosing regimes

Here we employ the previously-described fitting methodology to fit the mean normalized cell count of COLO858 cells subject to a fixed concentration of vemurafenib [32]. The doses of 0.032 *µ*M, 0.32 *µ*M, and 3.2 *µ*M were independently fit to the model in equations (16) - (17), and the results of our fitting can be seen in Figure 2(b)-(d). We observe an excellent agreement between the best-fit model solution and the three datasets. The optimal parameter values are provided in Table 1.

**Table 1:** Definition and best-fit value of parameters for the model in equations (16) - (17) over three drug concentrations. We also include the net resistant population growth rate *r*_*R*_ − *d*_*R*_, which represents the exponential decay rate of the resistant population in the absence of sensitive cells. Note that this net rate is most negative at the intermediate dose of 0.32 *µ*M.

| Parameter | Meaning | Drug Concentration |  |  |
| --- | --- | --- | --- | --- |
| | | 0.032 $\mu\text{M}$ | 0.32 $\mu\text{M}$ | 3.2 $\mu\text{M}$ |
| $r_S$ | Sensitive cell growth rate | 0.0532 | 0.0722 | 0.0880 |
| $d_S$ | Drug-induced sensitive cell death rate | 0.1855 | 0.4012 | 0.6924 |
| $\alpha$ | Rate of drug-induced resistance | 0.2520 | 0.1963 | 0.1048 |
| $r_R$ | Resistant cell growth rate | 0.0532 | 0.0415 | 0.0070 |
| $d_R$ | Drug-induced resistant cell death rate | 0.0751 | 0.0699 | 0.0137 |
| $r_R - d_R$ | Resistant cell net growth rate | -0.0219 | -0.0284 | -0.0067 |

Confidence in model parametrization is quantified in Fig. 3. As detailed in Section 3.2, model fitting is attempted for *M* = 1000 sets of initial parameter guesses. If realization *i* of the fitting algorithm results in cost function value *ζ*_*i*_ within 5% of the optimal cost function value *ζ*_*opt*_, the corresponding best-fit parameters are classified as suboptimal. Figure 3 displays the distribution of the suboptimal parameters for drug concentrations of 0.032 *µ*M, 0.32 *µ*M, and 3.2 *µ*M. We find that, for each model parameter, the suboptimal values are tightly distributed about the optimal value across the three doses, which can be interpreted as a measure of practical identifiability of model parameters; structural identifiability is justified in Section A.4. Further, each parameter behaves monotonically with respect to dose. We now discuss the behavior of each model parameter as a function of dose in detail.

The growth rate *r*_*S*_ of sensitive cellsis observed to monotonically increase as a function of drug dose, with no overlap in the suboptimal distributions across the three doses. This counterintuitive observation may be a consequence of a relatively fast cellular response to stress (chemotherapy), which prompts a release of growth signals via paracrine signaling; this has been observed in the case of pancreatic ductal adenocarcinoma upon radiotherapy [45]. Interestingly, this paracrine mechanism is mediated by the AKT/p38 and JNK pathways, which are typical bypass signaling pathways activated in melanoma following BRAF inhibitor treatment [31]. Other possible mechanisms might include one or more of the following when a targeted inhibitor such as vemurafenib is applied: (1) an increase in growth rate as the cell is freed from its regulated growth constraints; (2) activation of an alternate compensatory pathway that is more efficient, allowing energy or other resources being redirected toward growth; or (3) inhibition of cell cycle checkpoints that prevent division.

As expected, the drug-induced death rate of sensitive cells *d*_*S*_ also monotonically increases as a function of drug concentration. Again we find no overlap in the suboptimal distributions across the three doses, indicating that drug-induced cell kill behaves in a dose-dependent manner.

The rate of drug-induced resistance *α* is found to monotonically decrease as a function of drug concentration. As opposed to the sensitive population growth and induced death rates *r*_*S*_ and *d*_*S*_, a slight overlap exists in the distribution of *α* over the lower two doses. In particular, the maximal suboptimal value of *α* found at dose of 0.32 *µ*M is 0.2518, whereas the minimal value found at a dose of 0.032 *µ*M is 0.2461. This overlap is visually represented in Figure 3 by the absence of the hatch marks between the distributions. We had originally hypothesized that the relationship observed between *α* and drug dose would be the reverse of what is observed; that is, we had hypothesized that higher drug doses would induce resistance at a quicker rate. A possible explanation for this behavior is that higher dosages may kill the sensitive cell population (as a result of the increased value of *d*_*S*_) prior to the cell being able to transition to a resistant state.

The distributions of the resistant subpopulation parameters tell an interesting story about the behavior of drug-resistant cells that form at higher drug doses. Focusing first on the growth rate *r*_*R*_, we observe the expected relationship: this rate decreases as a function of drug dose. Interestingly, the order of magnitude of the suboptimal growth rates are the same at the two lower doses of the drug. In particular, the suboptimal values at the dose of 0.032 *µ*M are in the range [0.0498, 0.0546]. For the dose of 0.32 *µ*M, the range of suboptimal parameter is [0.0353, 0.0445]. Although a dose-dependence between these lower doses exists, the *r*_*R*_ parameter is always on the order 10^−2^. We observe a different phenomenon at the highest dose of 3.2 *µ*M, where more than 98% of the suboptimal parameters are on the order of 10^−3^. Similarly, suboptimal values of the drug-induced death rate *d*_*R*_ range from [0.0001, 0.0417] at the highest drug concentration, as compared to [0.0581, 0.0752] at the intermediate concentration and [0.0691, 0.0779] at the lowest concentration. Taken together, we find that resistant cells that were induced by the highest drug concentration grow, and are killed by the drug, at significantly lower rates than at intermediate and low drug concentrations. This result is worth comparing to a finding in [32] that COLO858 cells treated at a dose of 3.2 *µ*M exhibit one of the three responses: rapid death, quiescence (the cells were observed to survive, but not divide, in the 120 hour experimental time window), and resistant-like behavior where the surviving cells divide at much lower rates than observed in the absence of drug. The fact that, only at the highest dose in our model, the resistant subpopulation takes on quiescent-like behavior (very slow growth and death) seems consistent with the response to high drug doses in [32].

We also observe an interesting relationship between the *net* resistant growth rate, *r*_*R*_ − *d*_*R*_, and dose, which is provided in the last row of Table 1. As this rate is always negative, we see that all of the applied doses are able to asymptotically eliminate the resistant population. However, this net rate is most negative at the intermediate dose of 0.32 *µ*M, suggesting that intermediate dosing regimens may be more effective than a maximally tolerated dose (MTD) inspired strategy. This is investigated more thoroughly in Section 4.4, where we formulate an optimal control problem to determine the effectiveness of different treatment protocols. Note that *r*_*R*_ − *d*_*R*_ represents the asymptotic net growth rate of the resistant population, as the sensitive population is always eliminated (*r*_*S*_ − *d*_*S*_ is negative for all doses in Table 1).

Taken together, the tight distribution of the best fit parameter values across three drug concentrations, as well as the monotonicity of the parameter distributions as a function of drug concentration, lends support to the sufficiency of the two-compartment model proposed of drug-induced resistance in equations (16) - (17). We believe that these results, with its concomitant quantitative unraveling of the contributions of the two subpopulations as shown in Figure 2, reveal a biologically meaningful clustering of cells into just two major subclasses; we note that is this an experimentally testable hypothesis, as is briefly discussed in Section 6. However, it is reasonable to ask whether the resistant and quiescent cells should be modeled separately, given the three distinct responses to high drug concentrations observed in [32]. We explore this in the following section.

### 4.2 A finer-grained three-population model does not substantially improve fits to data

Given that experimental data indicates the presence of a subpopulation of quiescent cells at the highest drug dose that is distinct from the resistant subpopulation, we next explored whether a three population model that accounts for both a quiescent and a resistant subpopulation would better describe the experimental data. The three-population model we consider is a direct extension of our two-population model:

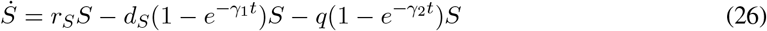

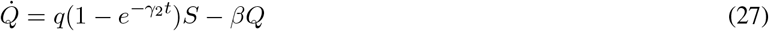

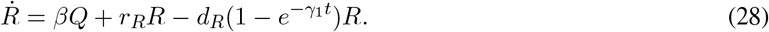

In this model, *Q* represents the normalized number of quiescent cells, and it is assumed that cells must pass through the quiescent state before becoming resistant. The drug-induced transition of sensitive cells to quiescent cells occurs at rate *q*, and quiescent cells transition to resistant at rate *β*. Other than these changes, the three-population model has the same assumptions as the two-population model, and all common parameters have the same definition as the two-population model. We again assume that *γ*_1_ = *γ*_2_ = 0.01, and that the initial population is composed of all sensitive cells, here meaning *S*(0) = 1, *Q*(0) = *R*(0) = 0.

The best fits of the three-population model to the experimental cell counts at a dose of 0.032, 0.32 and 3.2 *µ*M are shown in Figure 4(a)-(c). Although excellent fits to the data are obtained, the fits do not show any noticeable improvements from the two-population model (see Fig. 2). This visual observation is confirmed by comparing the optimal value of the cost function using the two- and three-population model (Figure 4(d)).

**Figure 4:**
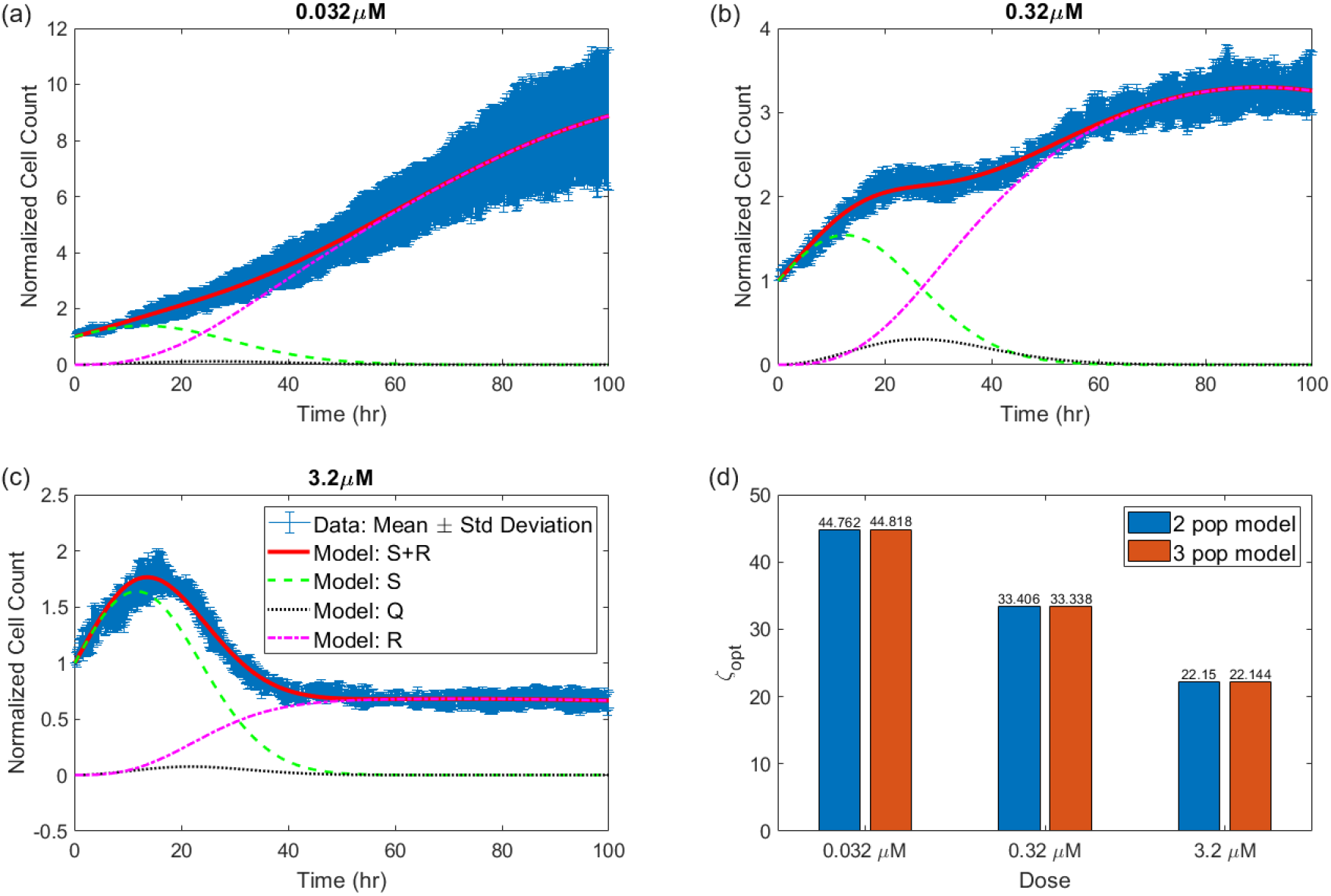
Model fits for the three-population model. Best model fits are shown at a drug concentration of (a) 0.032 *µ*M, (b) 0.32 *µ*M, and (c) 3.2 *µ*M. (d) The optimal value of the cost function, *ζ*_*opt*_ when data from each dose is fit using either the 2-population or the 3-population model.

Plots of the suboptimal parameter distributions for the three-population model (Fig. 5) further highlight the lack of necessity of a three-population model, as many of the model parameters are non-identifiable. The exception to this are the parameters describing the sensitive cells. Not only do *r*_*S*_ and *d*_*S*_ have tight distributions in the three-population, the distributions are incredibly similar to the two-population case in Fig. 3. The lack of identifiability is demonstrated quite clearly for the resistant cell parameters at the dose of 3.2 *µ*M. While the resistant cell growth rate *r*_*R*_ behaves quite similarly in the two- and three-population model at lower doses, at the dose of 3.2 *µ*M, the suboptimal parameters for the three-population model extend over a very wide range of values, indicating that we do not have sufficient data to identify the value of the parameter in this case. Interestingly, the drug-induced death rate *d*_*R*_ of resistant cells differs across doses when comparing the two- and three-population model. Though, just like with *r*_*R*_ the loss of identifiability is most apparent at the highest drug dose. Interestingly, the transition parameters *q* and *β* preserve their identifiability at higher drug doses, but lose identifiability at the lowest dose of 0.032 *µ*M. Taken together, this analysis indicates that a number of model parameters cannot be adequately identified given the available data when using the three-population model. The absence of improvement in model fits, coupled with the lack of identifiability of model parameters, lends strong support to the sufficiency of the two-population model.

**Figure 5:**
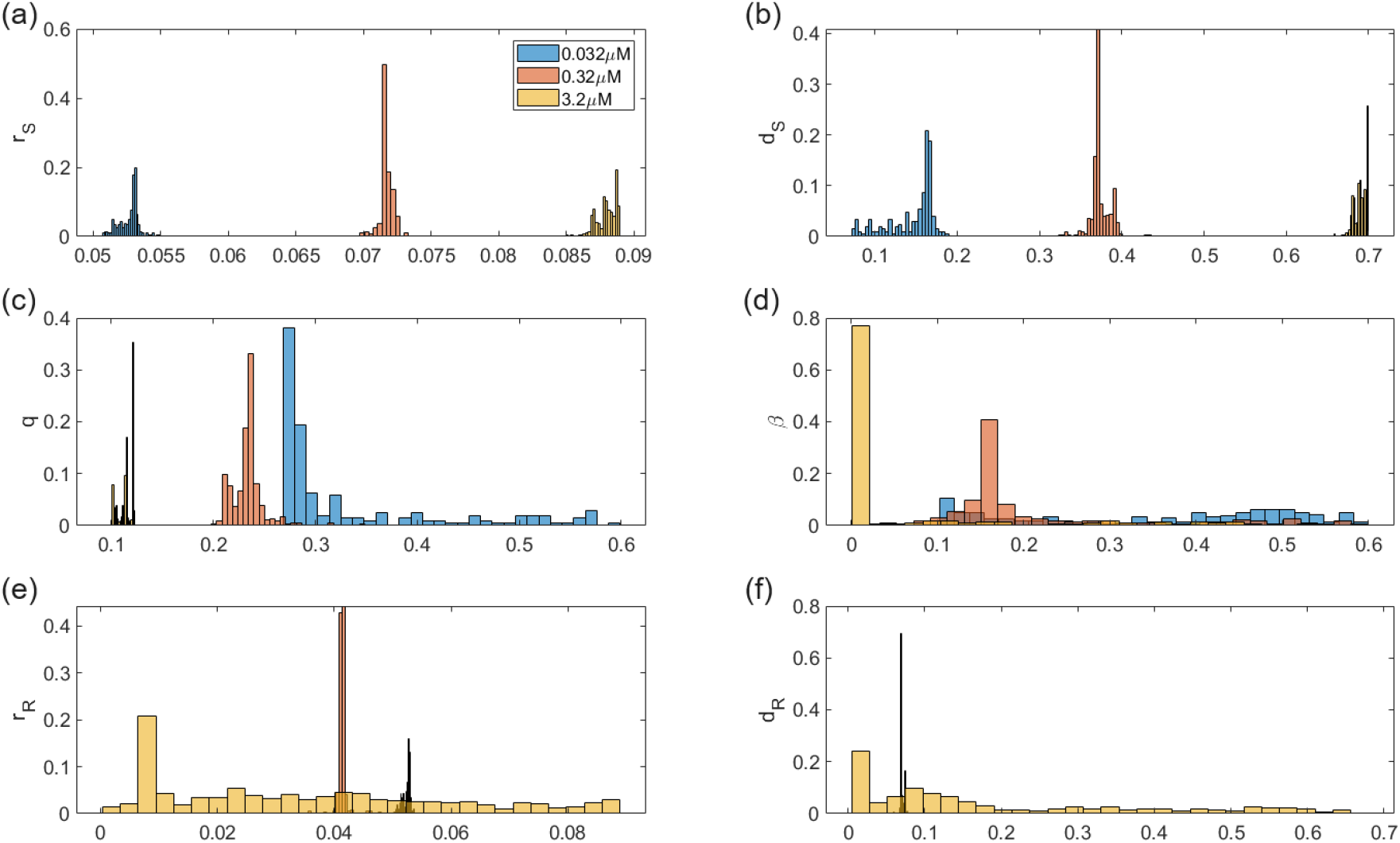
Practical identifiability of parameters in the three-population model. Distribution of parameters that give fits within 5% of the optimal parametrization for three-population model at three doses: 0.032 *µ*M in blue, 0.32 *µ*M in red, and 3.2 *µ*M in yellow. Plots are shown for (a) *r*_*S*_, (b) *d*_*S*_, (c) *q*, (d) *β*, (e) *r*_*R*_, and (f) *d*_*R*_.

### 4.3 Predicting population response at other doses

Given our choice to proceed with the two-population model, we next turned to validation of that model. Model fitting was performed for doses of 0.032 = 10^−1.5^*µ*M, 0.32 = 10^−0.5^*µ*M, and 3.2 = 10^0.5^*µ*M, with data at the doses of 0.1 = 10^−1^*µ*M and 1 = 10^0^*µ*M withheld for validation purposes. Observing that all experimental doses are uniformly spaced on a log 10 scale, we performed a piecewise linear interpolation of the best-fit parameters in Table 1 as a function of the log (base 10) of drug concentration. We used the interpolation to predict parameter values for log-dose values of {−1, 0}; that is, for doses of 0.1*µ*M and 1*µ*M. The interpolation process and resulting parameters are shown in Fig. A.1.

In Fig. 6, we illustrate the goodness-of-fit of the model solved at the interpolated parameter sets to the *in vitro* cell count data at the dose of 0.1 and 1 *µ*M. We observe that the model is able to well-capture the qualitative dynamics of tumor growth at the two doses that were withheld for fitting purposes. Quantitatively, the fits are quite good, though there are some notable shortcomings. At the 0.1 *µ*M dose, model predictions are at the upper bound of the experimental data. At the 1 *µ*M dose, the model predicts that the normalized cell counts achieves its maximum several hours earlier than is observed in the data, while also predicting local minimum around hour 40 that is not seen in the *in vitro* data. Given that parameters at both doses were obtained by a simple piecewise linear interpolation of parameter values obtained at other doses, the quality of these predictions does indicate that our model capable of describing COLO858 response to a wide range of vemurafenib doses.

**Figure 6:**
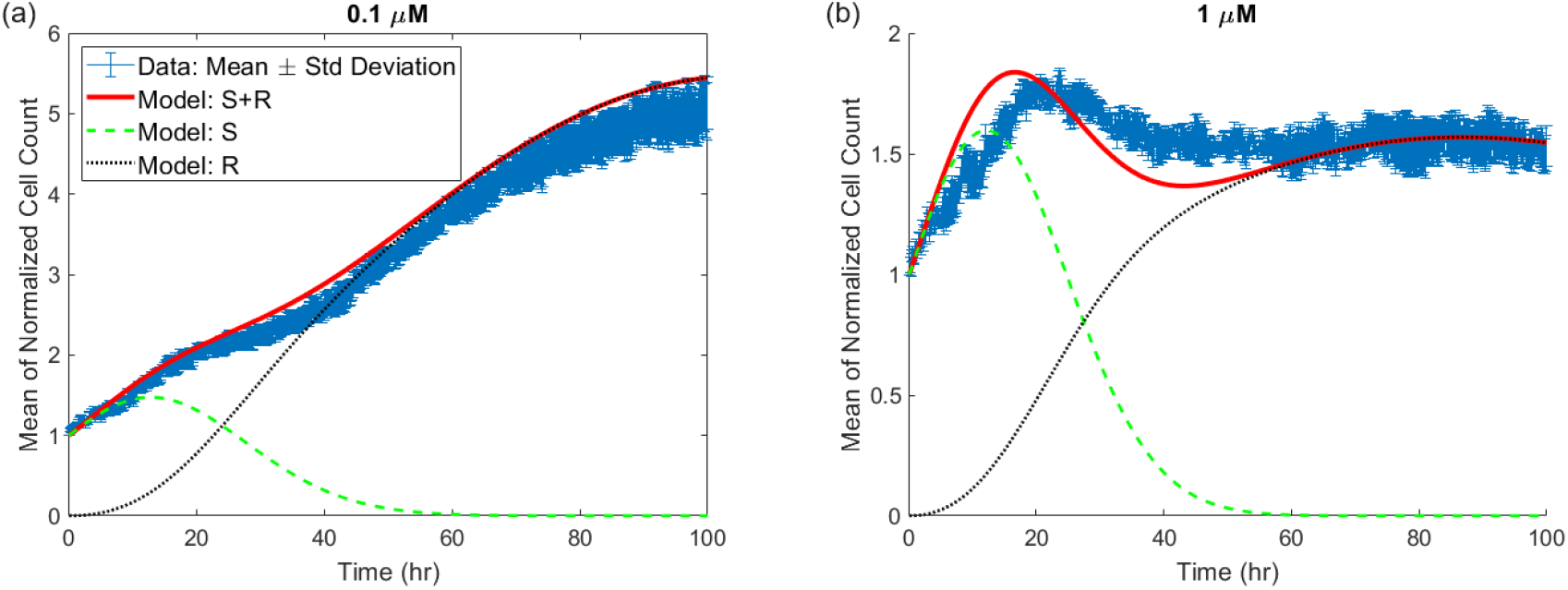
Data and model predictions using interpolated parameters. Predictions are shown at a drug concentration of (a) 0.1 *µ*M and (b) 1 *µ*M.

**Figure 7:**
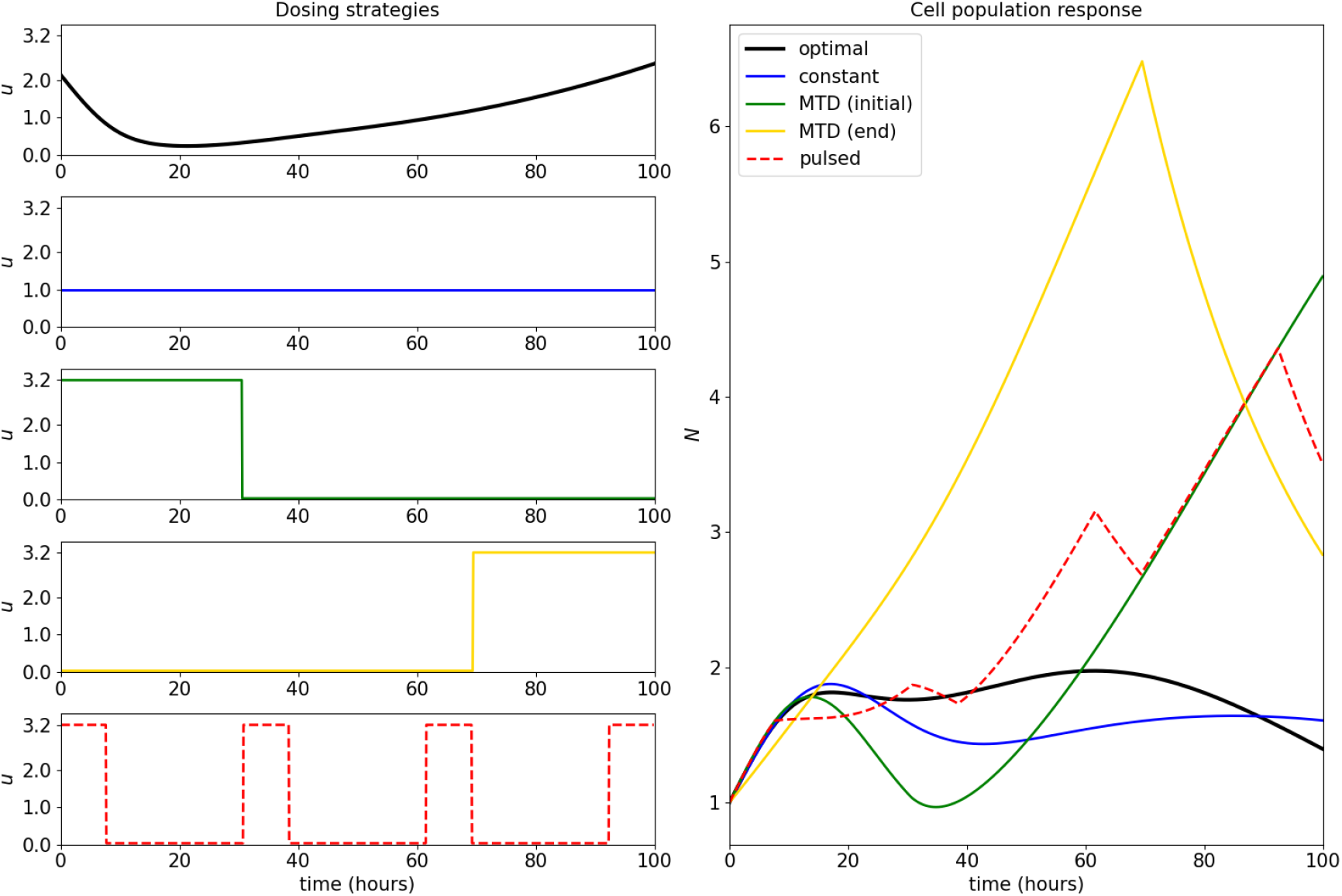
Response of cell population to applied controls. Control strategies considered appear on the left, and include the numerically computed optimal control (black) a constant intermediate dose (blue), a maximal tolerated dose applied at the onset of therapy (green), a maximal tolerated dose applied at the end of therapy (gold), and a pulsed “bang-bang” strategy with four “on” cycles (red, dotted). All strategies obtain the same total applied dosage of *M* = 100 *µ*M hours. Total cell population (*N* = *S* + *R*) responses are provided in the right figure. Note that the numeric optimal control yields a smaller final tumor size compared to the other strategies considered, but is comparable in value to the constant intermediate (i.e. metronomic) therapy.

### 4.4 Optimal control results

The parameterization of the two-compartment model in Table 1 suggests that high drug doses may not be optimal. While the induction rate *α* decreases as a function of dose, the drug-induced resistant cells are actually most-targetable at an intermediate drug dose. This can be seen in the values of resistant cell net growth rate inclusive of drug effect *r*_*R*_ − *d*_*R*_ in Table 1. The larger the value of *r*_*R*_ − *d*_*R*_ the “more resistant” the drug-induced phenotype is. As this combined parameter is minimized at the intermediate dose of 0.32 *µ*M, and the induced resistance rate is not maximal at this dose, together this indicates an intermediate dose of drug may be optimal.

As our mathematical model has been calibrated and validated on a numbers of doses, we can use it to determine theoretical optimal dosing strategies, as well as how close “standard” dosing protocols are to being optimal. Here, we explore solution to the optimal control problem posed in Section 3.3. In particular, we numerically solve (23) subject to (24) and (25) via CasADI, where we have constructed smooth approximations to the rate function *θ*(*u*) as discussed in Section A.3. Recall that we are interested in minimizing the cancer cell population *S* + *R* at a time *t*_*f*_ ; as our model is only validated on the experimental time window *t* ∈ [0, 100] hours, we fix *t*_*f*_ = 100 hours (see Figure 2, for example). We also assume an intermediate total applied dosage of *M* = 100 *µ*M hours, which corresponds to the constant dose of *u* ≡ 1 *µ*M applied throughout [0, *t*_*f*_ ]. The computed optimal control, as well as the population response, is provided in Figure 7. We also include a comparison with a number of other dosing strategies, including a constant intermediate dose, two maximally tolerated dose (MTD) strategies (one where the dose is applied at the beginning of the treatment, and the other where it is applied at the end), as well as a periodic “bang-bang” switching strategy. Note that all applied controls, including the optimal control, reach the upper bound on the total applied dosage *M*, and thus are all comparable with respect to the total amount of applied drug.

The structure of the computed optimal control confirms the prediction discussed in Section 4.1 regarding the optimality of intermediate doses: the optimal dosing regimen remains well below the upper bound of 3.2 *µ*M. Moreover, while this control may be difficult or even impossible to implement, we observe that it is well approximated by the constant intermediate dosing strategy *u*(*t*) ≡ 1 *µ*M. This is especially clear when we compare the final cancer cell populations, with the optimal control only slightly outperforming the constant strategy. As the constant therapy is an idealization of metronomic therapy (i.e. low dose, frequently or continuously administered chemotherapy [46]), our results imply that such therapy is near optimal. Comparing the population responses to the MTD therapies, we see that this is even more apparent, as this approximate metronomic therapy results in substantially smaller relative number of cancer cells (*N* (*t*_*f*_) ≈ 1.61, 4.89, 2.83 for constant, initial MTD, and ending MTD, respectively). For this cell population, it thus appears that MTD treatments are far from optimal, which is supported by a number of recent experimental and theoretical studies [47–53]. Furthermore, although the optimal control yields a slightly smaller final cancer size compared to constant therapy, it only achieves this result at the very end of the therapeutic window, and for most of the experiment, constant therapy results in a smaller cancer burden (e.g. the constant treatment has a smaller *L*^1^ norm).

## 5 Comparisons With Other Mathematical Models

As discussed in Section 1, the mathematical model used in this paper is a modification of the one that we had introduced in [33, 39]. In this section, we briefly review several deterministic alternative models of induced resistance that have been proposed in the recent literature.Although different, some of these models share important features with ours. See also the the review article [54].

The equations presented in the paper [55] model the dynamics of phenotypic adaptation in cell subpopulations, both on- and off-treatment, by a monoclonal antibody therapy (cetuximab, used on colorectal cancer cells). The equations during treatment are as follows:

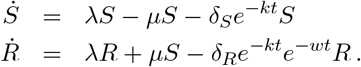

The cell proliferation rate *λ* is assumed to be the same in both sensitive (*S*) and resistant (*R*) populations. When the treatment is on, *S* cells may acquire resistance, and they switch to *R* cells at a rate *µ*. Cytotoxic effects on *S* and *R* are modeled by two linear terms, −*δ*_*S*_*e*^−*kt*^ and −*δ*_*R*_*e*^−*kt*^ respectively. The efficacy of the drug is assumed here to diminish at an exponential rate *e*^−*kt*^, because of the declining function of effector cells such as NK cells. The term *e*^−*wt*^ is used to model transience of resistance, which evolves under selective pressure at rate *w*. When the treatment is off, a different set of equations is used. In this regime, resistant *R* cells switch back to sensitive cells *S* at a rate of 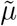, but no cell death is assumed:

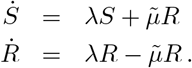

In contrast to our model, the rates of death of both the *S* and *R* subpopulations decrease with time (more for *R* than for *S* due to the term *e*^−*wt*^ representing transience of resistance), and the growth rates are the same for both subpopulations.

The paper [56] employs a simpler model of resistance to targeted therapy (particular, the EGFR inhibitor gefitinib) or classical chemotherapy (paclitaxel). The authors focus on the fact that, under therapy, tumor volume initially decreases, but eventually increases again due to resistance. There is one tumor volume (or total number of tumor cells) equation, which can also be found in [57]:

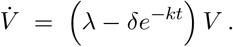

Here, *δ >* 0 quantifies the maximum drug effect, which occurs at the beginning of treatment. This effect exponentially decreases as resistance arises, and *λ >* 0 represents the net growth rate after the drugs have lost its effect due to resistance. It is assumed that *δ > λ*, so that at small times, prior to the emergence of resistance, the drugs are effective (negative net growth rate). Observe that, in contrast to our model and the model in [55], there is a single tumor volume equation instead of separate equations for sensitive and resistant cells. Interestingly, if we set *δ*_*R*_ = *δ*_*S*_ = *δ* and *w* = 0 (no transience of resistance) in the model in [55] then we recover these models from [56, 57] for the total cell population *V* = *S* + *R*.

The paper [58] employs a tumor growth model to evaluate patient-specific resistance to radiation combined with pembrolizumab and bevacizumab treatments for high-grade glioma. There are two equations, one for tumor growth dynamics and one for the death rate:

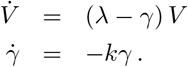

where *k* indicates the rate at which resistance develops. With *δ* = *γ*(0), this can be summarized by *dV/dt* = (*λ* − *δe*^−*kt*^)*V*, which is the same model as in [56, 57].

The paper [59] has a more complex model, involving three subpopulations: sensitive (*S*), persister (*P*), and resistant (*R*):

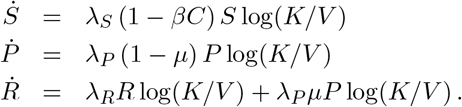

Here *K* is a carrying capacity term, *V* = *S* + *P* + *R, λ*_*i*_ are the Gompertz growth rate constants in the absence of drug, a proportionality term *βC* models death of sensitive tumor cells as a function of drug plasma concentration *C*(*t*), and the persister population *P* transitions to the resistant compartment with a mutation probability *µ*. This model differs from ours in many ways, including the Gompertz growth term (we tested Gompertz models and obtained poorer fits to our data), the lack of delays for drug action, and the use of three compartments, which we found did not improve fits.

The paper [60] focuses on melanoma adaptive therapy through two ODE models of interactions between drug-sensitive and resistant cells. In a first model, drug-sensitive (*S*) and drug-resistant (*R*) subpopulations compete for limited resources through a classical Lotka-Volterra competition for resources mechanism:

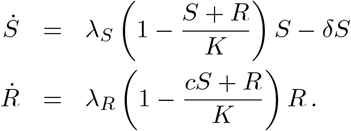

Here *K* is a carrying capacity and the parameter *c* quantifies the degree to which sensitive cells inhibit the growth rate of resistant cells in comparison to resistance cells among themselves. The authors assume that *λ*_*S*_ = 0 during treatment, because they model BRAF/MEK inhibitors which act through growth inhibition as well as apoptosis (*δ* term). An alternative model proposed in the same paper sets *c* = 1 but incorporates phenotypic switching between drug-sensitive and drug-resistant cells, with induced transitions from sensitive to resistant states (term *αS*) or vice-versa (term *βR*):

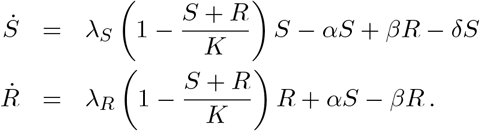

It is assumed that the transition rate constant *α* is non-zero during treatment, and is zero otherwise, and that the transition rate constant *β* is nonzero when treatment is off and is zero otherwise. This model is similar in many ways to our model in [33], which is cited in [60], but with several important differences, including the fact that the population-switching terms are only active during therapy or not, and that *R* cells are fully resistant (no death term).

The paper [61] developed a model for induced resistance of colorectal cancer cells to targeted therapies. The model has sensitive (*S*) and drug-tolerant persister (*P*) populations:

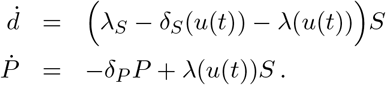

Drug-treated cells can die, replicate, or switch to a persister state *P* at a rate *λ* in a dose-dependent fashion *λ*(*u*(*t*)). *u*(*t*) is the amount of drug at time *t*, and it is either constant or linearly increasing (see [61], Supplementary Information, Extended Data Figure 5). Resistant cells are not explicitly modeled, because it is assumed that persisters that attempt to divide before acquiring drug-resistance mutations die, and thus a possible back-switching from persister to sensitive populations in the presence of drug would simply be incorporated into their death rate. Note that the death rate *δ*_*S*_(*u*(*t*)) of sensitive cells is assumed to be drug-dependent, but that of persister cells is not. This model differs from ours in many ways, including the fact that we have delays in drug action and phenotype transitions, and that a reverse phenotype transition back to sensitive type is non-negligible (and needed for fits). Note, however, that our three-population equations with a “quiescent” population could be thought of as having a persister compartment.

## 6 Conclusions

Drug-induced resistance is a major impediment to the success of targeted therapy. Yet, there are few mathematical models that are both simple and give excellent fits to tumor cell count or volume data when resistance is known to be induced by the drug. This paper describes a minimal mathematical model of drug-induced resistance and shows that it is able to provide excellent fits to time-resolved *in vitro* experimental data. Moreover, using only such bulk data on total cell numbers, the model allows one to separate the contributions of sensitive and resistance subpopulations and describes their dynamics. A theoretical identifiability analysis demonstrated that the parameters of the proposed model are, in principle, uniquely obtainable from data. A practical identifiability analysis showed tight confidence bounds on all parameters given the available data, provided we only consider a sensitive and resistant subpopulation of cells. Practical identifiability is lost in a finer-grained model that also includes a quiescent subpopulation. To determine the predictive power of the model, we assessed its ability to predict data at two untrained drug doses. Excellent qualitative, and reasonable quantitative, predictions were made by the model at both untrained doses. Using the validated model, we numerically explored an optimal control problem, with the goal of minimizing the final cell count subject to an applied dosage constraint. While we did identify a non-standard dosing strategy as the optimal control, the optimal control only slightly outperformed a metronomic-like protocol of administering a constant, lower dose of drug. On the other hand, maximal tolerated dose-like protocols proved to be far inferior.

This study suggests a variety experimental follow-ups. One of the most important of these is the validation of our predictions, at least qualitatively if not quantitatively, of the respective time-resolved contributions of resistant and sensitive populations. Our mathematically derived *S* and *R* populations suggest the possibility to group the distributions of mRNA, protein, or epigenetic dynamic signatures of individual cells into just two major clusters with separate drug sensitivity and growth and death rates. These signatures might reflect resistance markers, such as the increased expression of membrane transporters that enhance the efflux of drugs as in the study [21]. Experimental tools that could be utilized to investigate this include the use of single cell clonally-resolved transcriptome datasets (scRNA-seq) as in [62].

Another direction for work is the comparison of drug dosing strategies, for the same *in vitro* cell lines and drugs, as investigated in the mathematically derived optimal control analysis (Section 4.4). Many natural questions exist. For example, is a continuous-dose or a constant “metronomic” strategy actually superior to an MTD approach? Are other periodic strategies superior in terms of the final population size at a specific time horizon? Do pulsed strategies not afford any advantages, as with our example (red dashed curve) in Figure 7?

We speculated that the increase in growth rate of the sensitive tumor population may be due to its intrinsic response to stress, which happens at a faster time scale than the emergence of the resistant population arising from slower mechanisms such as epigenetic reprogramming. A deeper study of this issue is necessary. Is this a true effect, due to different time scales for different forms of resistance, with faster time scale signaling processes being reflected in *r*_*S*_ and slower gene expression or even epigenetic modifications leading to an increase in the delayed death rate *d*_*S*_? Identifying such mechanisms and then blocking their action may elucidate the relation between these two dose-dependent effects.

Of course, the development of similar models for other rich time-resolved data sources would also be of interest. Different cell lines and different drug choices will affect the parameters differently, through interference with metabolic processes or disruption of regulatory feedback loops.

## 7 Acknowledgements

The authors acknowledge Dr. Mohammad Fallahi-Sichani for his helpful discussions and access to the data. EDS acknowledges support from grants AFOSR FA9550-21-1-0289 and NSF/DMS-2052455. JLG thanks Aahna Rathod and Swetha Yogeswaran for their efforts related to this project using a different dataset. JMG thanks Arthur Castello Branco de Oliveira for his helpful discussions on implementing optimal control problems in CasADI.

## A Appendix

### A.1 Setting for fmincon

These are the settings used in the fmincon MATLAB^®^ code.

**Table A.1:** Settings used in MATLAB’s implementation of *fmincon*.

| Setting | Value |
| --- | --- |
| MaxFunctionEvaluations | 30,000 |
| Lower bound: $r_S$ | 0.001 |
| Lower bound: all other parameters | $10^{-6}$ |
| Upper bound: $r_S$ | 0.15 |
| Lower bound: all other parameters | 0.6 |

### A.2 Parameter interpolation

**Figure A.1:**
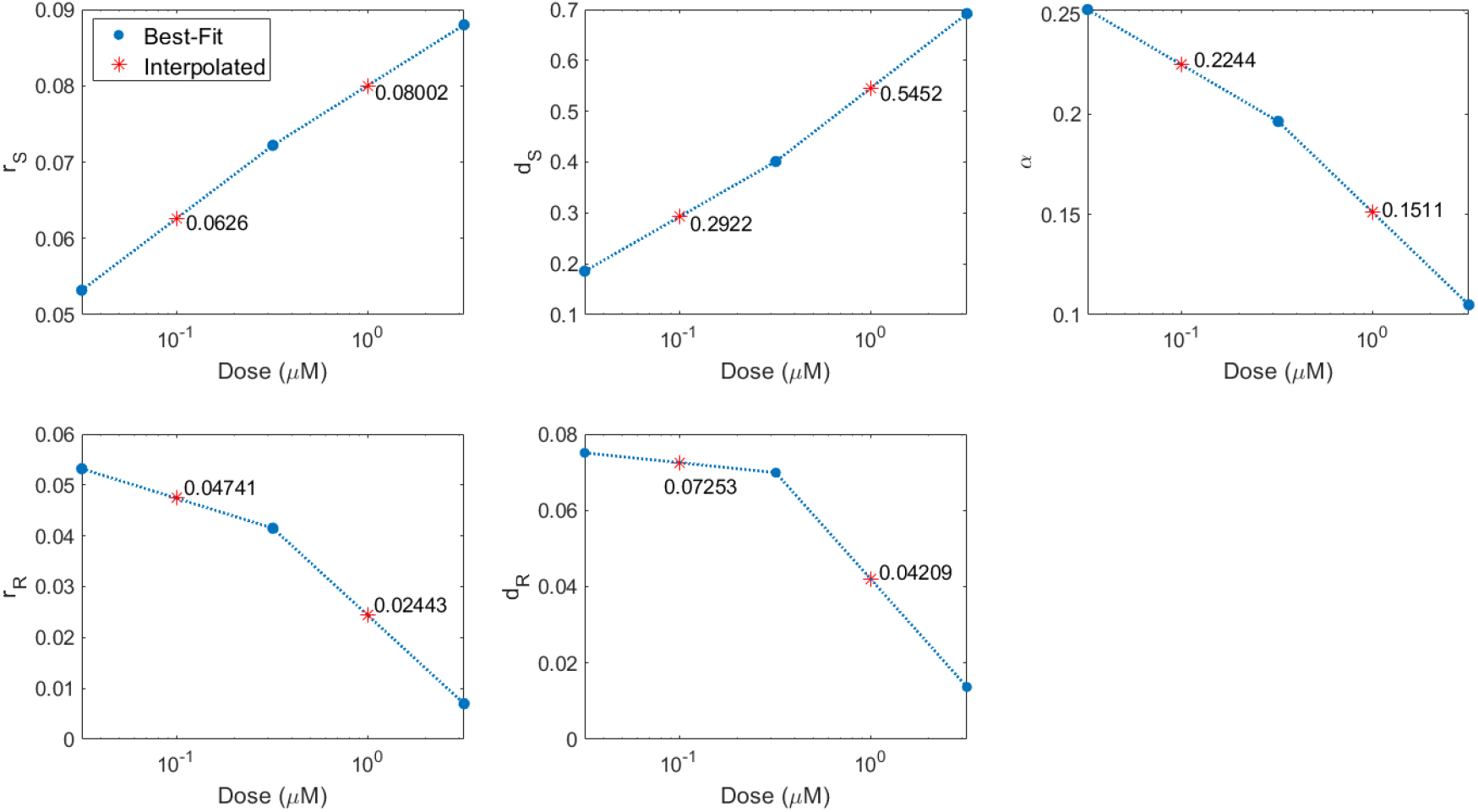
Parameter predictions. Piecewise linear interpolation (dashed blue line) of best-fit parameter values at drug concentrations 0.032, 0.32 and 3.2 *µ*M to predict parameter value at withheld concentrations of 0.1 and 1 *µ*M.

### A.3 Rate approximation

We have fit the two-compartment model (16)-(17) to obtain parameter values as a function of dose; see Table 1 for the estimated parameters *θ* = *θ*(*u*), where

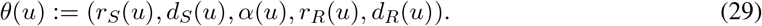

Recall that the drug-induced delay terms (*γ*_1,2_) were not fit, but fixed at a value of 0.01 per hour. Our work in Sections 4.1 and 4.3 suggests that linearly interpolating the rate parameters *θ* with respect to the logarithm (base 10) of the dose provides close agreement to the experimental data. Thus, utilizing such an interpolating, we can obtain expressions for the rate parameters for *all* doses between 0.032 *µ*M and 3.2 *µ*M:

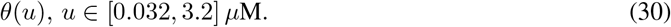

This interpolation is shown in Figure A.1.

As our objective in this section is to formulate and solve an optimal control problem, it will be numerically advantageous to assume that the vector field is smooth. More specifically, we will utilize CasADI [63], an open-source framework for nonlinear optimization and optimal control, which relies on the Interior Point Optimizer (Ipopt) software package to solve nonlinear programming problems. Here multiple shooting methods are utilized to transform the original optimal control problem to a constrained nonlinear programming problem, which is then solved via Ipopt. As Ipopt implements an interior line search method, it assumes that the objective is at least twice continuously differentiable [64], which is generally not satisfied by the function *θ*(*u*) in Figure A.1; recall *θ*(*u*) is derived from piecewise-linear interpolation on the logarithm of the doses. Thus, we approximate the piecewise-linear rate functions by degree three polynomials using standard least-squares regression; the original and approximating expressions are provided in Figure A.2. As the polynomial approximations closely match the piecewise-linear curves (in both the logarithmic and linear scales), for the remainder of the manuscript, we define *θ*(*u*) by the polynomial approximations in Figure A.2.

### A.4 Structural identifiability analysis

Our model (16)-(17), with fixed *γ*_1_ = *γ*_2_, is locally structurally identifiable in the following sense. Consider the set P of all possible parameters *θ* of the form

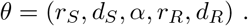

We assume that the initial conditions are *S*(0) = 1, *R*(0) = 0 (normalized cell counts). Local structural identifiability is a “well-posedness” property: it asks if different (but close by) parameters *θ*_1_ and *θ*_2_ should give different total cell counts *S*(*t*) + *R*(*t*). In other words, from (perfect) time-resolved measurements of *S*(*t*) + *R*(*t*), one should in theory be able to recover all parameters in a neighborhood of a given one. Structural identifiability analysis deals with the ideal case in which there are no measurement errors or replication variability. Although this is not a realistic assumption in practice, the goal of this analysis is to determine if there are parameter combinations that are redundant, in which case numerical fitting methods would not provide interpretable parameters due to non-uniqueness.

In the field of structural identifiability, one often looks for generic results. (Genericity is used in order to ignore accidental equalities. For example, if we are given the expression *px* and we with to determine *x*, this can always be done unless *p* happens to be zero, which is a very special parameter value.) Mathematically, we use the terminology in [65]: genericity means that there exists a “Zariski open” subset 𝒫_0_ of 𝒫 so that the identifiability property holds for pairs of vectors *θ*_1_ and *θ*_2_ in 𝒫_0_. A Zariski open set is defined by the non-vanishing of a nontrivial polynomial. Such a set is “generic” in the sense that its complement has (Lebesgue) measure zero, so with “probability 1” a randomly chosen set of parameters will be identifiable. We assume that *γ*_1_ = *γ*_2_ = *γ* is fixed and known. For showing identifiability, we can then take without loss of generality *γ* = 1 (the general case can be reduced to this one by a time-reparametrization, multiplying all parameters by the known *γ*). To simplify, we introduce new parameters

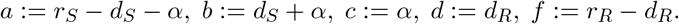

**Figure A.2:**
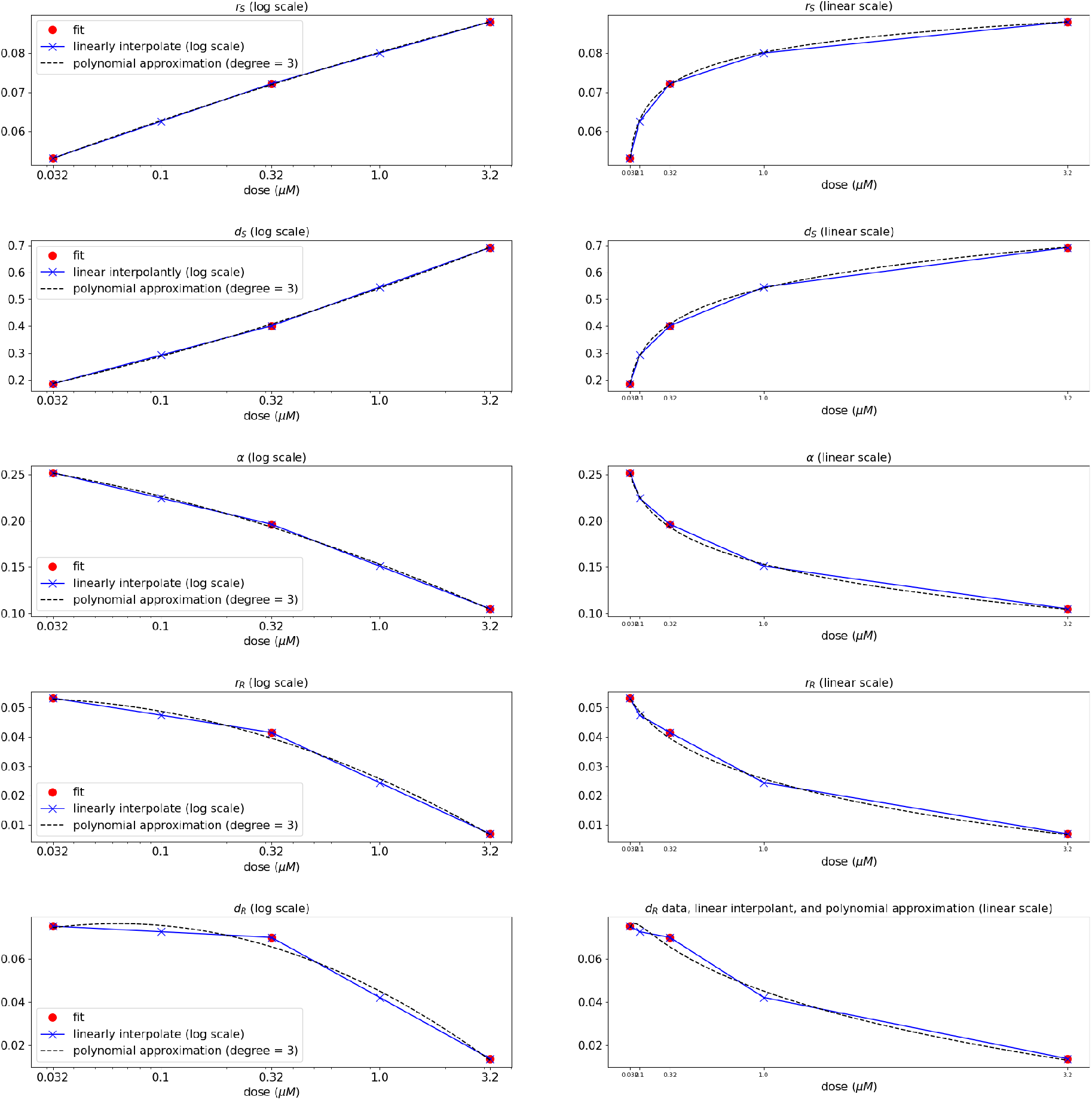
Rate parameters *r*_*S*_, *d*_*S*_, *α, r*_*R*_, and *d*_*R*_ in system (16) - (17) as a function of dose *u*. Red circles denote fits from experimental data, blue crosses denotes piecewise-linear interpolation (on the log scale with respect to dose), and the dotted black curve denotes a third-degree polynomial approximation to the data. The left column represents each rate parameter as a function of the logarithm of the dose, while the right provides the same relationship with dose scaled linearly. Compare to Figure A.1.

Identifiabilty with these new parameters is equivalent to the original ones, since the two sets are related by an invertible linear transformation. Our equations (16)-(17) become, with these notations:

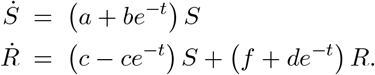

We view *y*(*t*) = *S*(*t*) + *R*(*t*) as the “measured output” of this system and compute the first five derivatives at time *t* = 0: *y*^′^(0) to *y*^(5)^(0). These derivatives are polynomial functions of the parameters. These computations can be done in closed form, using the product rule. For example, *S*^′^(0) = *a* + *b* and *R*^′^(0) = 0 (recall that *S*(0) = 1 and *R*(0) = 0). Similarly, *S*^′′^(*t*) = [(*a* + *be*^−*t*^)*S*(*t*)]^′^ = −*be*^−*t*^*S*(*t*) + [(*a* + *be*^−*t*^)]*S*^′^(*t*) = −*be*^−*t*^*S*(*t*)+[(*a*+*be*^−*t*^)]^2^, and hence *S*^′′^(0) = −*b*+(*a*+*b*)^2^. One has *y*^′^(0) = *a*+*b, y*”(0) = *c*−*b*−(*d*+*f*)(*b*− (*a*+*b*)^2^)+(*a*+*b*)^2^, and so forth. Let us call *F* the mapping from the 5-vectors of parameters (*a, b, c, f, e*) into the 5-vector of output derivatives at time 0, (*y*^′^(0), *y*”(0), *y*^(3)^(0), *y*^(4)^(0), *y*^(5)^(0)). The Jacobian of *J* is a 5× 5 matrix of parameters which will be nonsingular except at those parameter vectors where its determinant *D* is zero. Since *D* is a polynomial function of the parameters, nonsingularity holds on the Zariski open subset consisting of parameters for which *D*(*a, b, c, d, f*) is nonzero. The Implicit Mapping theorem then guarantees that this mapping is locally invertible around such points, meaning that one has at least identifiability in a local sense (any distinct but two close enough parameters give distinct outputs). It remains to actually verify that *D* is a nonzero polynomial. This can be done explicitly, For example when evaluated at *a* = 2 and *b* = *c* = *d* = *f* = 0, one computes *D*(2, 0, 0, 0, 0) = 576, which is nonzero. (Computations not shown.) This completes the proof of local generic identifiability.

It is also possible go further, and conclude global generic identifiability as well. To do this, we may apply a computational package such as the Structural Identifiability Toolbox (SIAN) [66]. In order to apply this software, we need to transform our system into one given by a polynomial system of equations. Thus we study instead the following system:

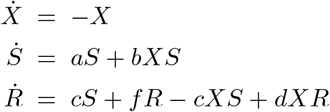

with outputs *S* + *R* and *X*. The software package SIAN (run online^1^) confirms global (generic) identifiability of this system, even when initial states are unknown and need to be identified simultaneously with the parameters. We also verified the local identifiability property using an alternative package, STRIKE-GOLDD [67], confirming the result.

It is interesting to note that the delay terms are essential for structural identifiability. If *γ* = 0 (no delay), the equations would become:

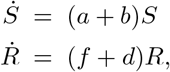

and is in that case impossible to distinguish between pairs (*a, b*) with the same sum *a* + *b*, nor pairs (*d, f*) with the same *d* + *f*

## Footnotes

1 available online at https://maple.cloud/app/6509768948056064

## References

[1] S. B. Zaman, M. A. Hussain, R. Nye, V. Mehta, K. T. Mamun, and N. Hossain, “A review on antibiotic resistance: alarm bells are ringing,” Cureus, vol. 9, p. e1403, Jun 2017.

[2] L. Menendez-Arias and D. D. Richman, “Editorial overview: antivirals and resistance: advances and challenges ahead,” Curr Opin Virol, vol. 8, pp. iv–vii, Oct 2014.

[3] J. Beardsley, C. L. Halliday, S. C. Chen, and T. C. Sorrell, “Responding to the emergence of antifungal drug resistance: perspectives from the bench and the bedside,” Future Microbiol, vol. 13, pp. 1175–1191, 08 2018.

[4] H. Zahreddine and K. Borden, “Mechanisms and insights into drug resistance in cancer,” Frontiers in Pharmacology, vol. 4, p. 28, 2013.

[5] S. Kim, T. M. Kim, D.-W. Kim, H. Go, B. Keam, S.-H. Lee, J.-L. Ku, D. H. Chung, and D. S. Heo, “Heterogeneity of genetic changes associated with acquired crizotinib resistance in alk-rearranged lung cancer,” Journal of Thoracic Oncology, vol. 8, no. 4, pp. 415–422, 2013.

[6] C. Holohan, S. Van Schaeybroeck, D. B. Longley, and P. G. Johnston, “Cancer drug resistance: an evolving paradigm,” Nature Reviews Cancer, vol. 13, no. 10, pp. 714–726, 2013.

[7] M. Gottesman, “Mechanisms of cancer drug resistance,” Annual Review of Medicine, vol. 53, pp. 615–627, 2002.

[8] M. Dean, T. Fojo, and S. Bates, “Tumour stem cells and drug resistance,” Nature Reviews Cancer, vol. 5, pp. 275–284, 2005.

[9] D. Woods and J. Turchi, “Chemotherapy induced DNA damage response,” Cancer Biology & Therapy, vol. 14, pp. 379–389, 2013.

[10] T. Gajewski, M. Y. C. Blank, I. Brown, K. A. J. Kline, and H. H., “Immune resistance orchestrated by the tumor microenvironment,” Immunological Reviews, vol. 213, pp. 131–145, 2006.

[11] O. Trédan, C. Galmarini, and I. Tannock, “Drug resistance and the solid tumor microenvironment,” Journal of the National Cancer Institute, vol. 99, pp. 1441–1454, 2007.

[12] M. Meads, R. Gatenby, and W. Dalton, “Environment-mediated drug resistance: a major contributor to minimal residual disease,” Nature Reviews Cancer, vol. 9, pp. 665–674, 2009.

[13] A. Correia and M. Bissell, “The tumor microenvironment is a dominant force in multidrug resistance,” Drug Resistance Updates, vol. 15, pp. 39–49, 2012.

[14] D. McMillin, J. Negri, and C. Mitsiades, “The role of tumour-stromal interactions in modifying drug response: challenges and opportunities,” Nature Reviews Drug Discovery, vol. 12, pp. 217–228, 2013.

[15] J. Foo, D. Basanta, R. Rockne, C. Strelez, C. Shah, K. Ghaffarian, S. Mumenthaler, K. Mitchell, J. Lathia, D. Frankhouser, S. Branciamore, Y. Kuo, G. Marcucci, R. Velde, A. Marusyk, S. Huang, K. Hari, M. Jolly, H. Hatzikirou, K. Poels, M. Spilker, B. Shtylla, M. Robertson-Tessi, and A. Anderson, “Roadmap on plasticity and epigenetics in cancer,” Phys Biol., vol. 19, no. 3, p. 031501, 2022.

[16] M. O. Sommer, G. M. Church, and G. Dantas, “The human microbiome harbors a diverse reservoir of antibiotic resistance genes,” Virulence, vol. 1, no. 4, pp. 299–303, 2010.

[17] Z. Iqbal, A. Aleem, M. Iqbal, M. Naqvi, A. Gill, and A. e. a. Taj, “Sensitive detection of preexisting BCR-ABL kinase domain mutations in CD34+ cells of newly diagnosed chronic-phase chronic myeloid leukemia patients is associated with imatinib resistance: implications in the post-imatinib era,” PloS ONE, vol. 8, p. e55717, 2013.

[18] C. Roche-Lestienne and C. Preudhomme, “Mutations in the ABL kinase domain pre-exist the onset of imatinib treatment,” Semin. Hematol., vol. 40, pp. 80–82, 2003.

[19] M. Carlino, C. Fung, H. Shahheydari, J. Todd, S. Boyd, M. Irvine, A. Nagrial, R. Scolyer, R. Kefford, G. Long, and H. Rizos, “Preexisting MEK1P124 mutations diminish response to BRAF inhibitors in metastatic melanoma patients,” Clinical Cancer Research, vol. 21, pp. 98–105, 2015.

[20] S. Luria and M. Delbrück, “Mutations of bacteria from virus sensitivity to virus resistance,” Genetics, vol. 28, pp. 491–511, 1943.

[21] A. O. Pisco, A. Brock, J. Zhou, A. Moor, M. Mojtahedi, D. Jackson, and S. Huang, “Non-darwinian dynamics in therapy-induced cancer drug resistance,” Nature communications, vol. 4, 2013.

[22] J. Nyce, S. Leonard, D. Canupp, S. Schulz, and S. Wong, “Epigenetic mechanisms of drug resistance: drug-induced DNA hypermethylation and drug resistance,” Proceedings of the National Academy of Sciences, vol. 90, pp. 2960–2964, 1993.

[23] J. Nyce, “Drug-induced DNA hypermethylation: a potential mediator of acquired drug resistance during cancer chemotherapy,” Mutation Research, vol. 386, pp. 153–161, 1997.

[24] S. Sharma, D. Lee, B. Li, and M. e. a. Quinlan, “A chromatin-mediated reversible drug-tolerant state in cancer cell subpopulations,” Cell, vol. 141, pp. 69–80, 2010.

[25] A. Pisco and S. Huang, “Non-genetic cancer cell plasticity and therapy-induced stemness in tumour relapse: ‘What does not kill me strengthens me’,” British Journal of Cancer, vol. 112, pp. 1725–1732, 2015.

[26] S. M. Shaffer, M. C. Dunagin, S. R. Torborg, E. A. Torre, B. Emert, C. Krepler, M. Beqiri, K. Sproesser, P. A. Brafford, M. Xiao, et al., “Rare cell variability and drug-induced reprogramming as a mode of cancer drug resistance,” Nature, vol. 546, no. 7658, p. 431, 2017.

[27] G. Housman, S. Byler, S. Heerboth, K. Lapinska, M. Longacre, N. Snyder, and S. Sarkar, “Drug resistance in cancer: an overview,” Cancers, vol. 6, pp. 1769–1792, 2014.

[28] P. K. Dhanyamraju, T. D. Schell, S. Amin, and G. P. Robertson, “Drug-Tolerant Persister Cells in Cancer Therapy Resistance,” Cancer Research, vol. 82, pp. 2503–2514, 07 2022.

[29] S. Ramisetty, A. R. Subbalakshmi, S. Pareek, T. Mirzapoiazova, D. Do, D. Prabhakar, E. Pisick, S. Shrestha, S. Achuthan, S. Bhattacharya, J. Malhotra, A. Mohanty, S. S. Singhal, R. Salgia, and P. Kulkarni, “Leveraging cancer phenotypic plasticity for novel treatment strategies,” Journal of Clinical Medicine, vol. 13, no. 11, 2024.

[30] K. Leder, J. Foo, B. Skaggs, M. Gorre, C. Sawyers, and F. Michor, “Fitness corrected by BCR-ABL kinase domain mutations determines the risk of pre-existing resistance in chronic myeloid leukemia,” PLoS ONE, vol. 6, p. e27682, 2011.

[31] M. Fallahi-Sichani, V. Becker, B. Izar, G. Baker, J. Lin, S. Boswell, P. Shah, A. Rotem, L. Garraway, and P. Sorger, “Adaptive resistance of melanoma cells to RAF inhibition via reversible induction of a slowly dividing de-differentiated state,” Mol Syst Biol., vol. 13, no. 1, p. 905, 2017.

[32] N. Comandante-Lou, M. Khaliq, D. Venkat, M. Manikkam, and M. Fallahi-Sichani, “Phenotype-based probabilistic analysis of heterogeneous responses to cancer drugs and their combination efficacy,” PLoS Comput Biol, vol. 16, p. e1007688, Feb 2020.

[33] J. Greene, J. Gevertz, and E. Sontag, “Mathematical approach to differentiate spontaneous and induced evolution to drug resistance during cancer treatment,” JCO Clin. Cancer Inform., vol. 3, pp. 1–20, 2019.

[34] I. Proietti, N. Skroza, S. Michelini, A. Mambrin, V. Balduzzi, N. Bernadini, A. Marchesiello, E. Tolino, S. Volpe, P. Maddalena, M. Di Fraia, G. Mangino, G. Romeo, and C. Potenza, “Braf inhibitors: Molecular targeting immunomodulatory actions,” Cancers, vol. 12, Jul 2020.

[35] S. M. Shaffer, M. C. Dunagin, S. R. Torborg, E. A. Torre, B. Emert, C. Krepler, M. Beqiri, K. Sproesser, P. A. Brafford, M. Xiao, E. Eggan, I. N. Anastopoulos, C. A. Vargas-Garcia, A. Singh, K. L. Nathanson, M. Herlyn, and A. Raj, “Rare cell variability and drug-induced reprogramming as a mode of cancer drug resistance,” Nature, vol. 546, pp. 431–435, Jun 2017.

[36] S. E. Shackney, G. W. McCormack, and G. J. Cuchural, “Growth rate patterns of solid tumors and their relation to responsiveness to therapy: an analytical review,” Annals of Internal Medicine, vol. 89, no. 1, pp. 107–121, 1978.

[37] W.-P. Lee, “The role of reduced growth rate in the development of drug resistance of hob1 lymphoma cells to vincristine,” Cancer Letters, vol. 73, no. 2-3, pp. 105–111, 1993.

[38] M. Berenbaum, “In vivo determination of the fractional kill of human tumor cells by chemotherapeutic agents,” Cancer chemotherapy reports, vol. 56, no. 5, pp. 563–571, 1972.

[39] J. Greene, C. Sanchez-Tapia, and E. Sontag, “Mathematical details on a cancer resistance model,” Frontiers in Bioengineering and Biotechnology, vol. 8, pp. 501: 1–27, 2020.

[40] E. Sontag, Mathematical Control Theory. Deterministic Finite-Dimensional Systems, vol. 6 of Texts in Applied Mathematics. New York: Springer-Verlag, second ed., 1998.

[41] F. Fröhlich, L. Gerosa, J. Muhlich, and P. K. Sorger, “Mechanistic model of MAPK signaling reveals how allostery and rewiring contribute to drug resistance,” Mol Syst Biol, vol. 19, p. e10988, Feb 2023.

[42] D. Beck, H. Niessner, K. S. Smalley, K. Flaherty, K. H. Paraiso, C. Busch, T. Sinnberg, S. Vasseur, J. L. Iovanna, S. en, B. Stork, S. Wesselborg, M. Schaller, T. Biedermann, J. Bauer, K. Lasithiotakis, B. Weide, J. Eberle, B. Schittek, D. Schadendorf, C. Garbe, D. Kulms, and F. Meier, “Vemurafenib potently induces endoplasmic reticulum stress-mediated apoptosis in BRAFV600E melanoma cells,” Sci Signal, vol. 6, p. ra7, Jan 2013.

[43] S. Kucherenko, D. Albrecht, and A. Saltelli, “Exploring multi-dimensional spaces: a comparison of latin hypercube and quasi monte carlo sampling techniques,” arXiv, no. 1505.02350, 2015.

[44] A. Bressan and B. Piccoli, Introduction to the mathematical theory of control, vol. 1. American institute of mathematical sciences Springfield, 2007.

[45] J. Cheng, L. Tian, J. Ma, Y. Gong, Z. Zhang, Z. Chen, B. Xu, H. Xiong, C. Li, and Q. Huang, “Dying tumor cells stimulate proliferation of living tumor cells via caspase-dependent protein kinase Cδ activation in pancreatic ductal adenocarcinoma,” Molecular oncology, vol. 9, p. 105—114, January 2015.

[46] N. C. Institute, “Definition of metronomic chemotherapy,” 2024.

[47] O. G. Scharovsky, L. E. Mainetti, and V. R. Rozados, “Metronomic chemotherapy: changing the paradigm that more is better,” Current oncology, vol. 16, no. 2, p. 7, 2009.

[48] E. Pasquier, M. Kavallaris, and N. André, “Metronomic chemotherapy: new rationale for new directions,” Nature reviews Clinical oncology, vol. 7, no. 8, pp. 455–465, 2010.

[49] Y.-P. Chen, X. Liu, Q. Zhou, K.-Y. Yang, F. Jin, X.-D. Zhu, M. Shi, G.-Q. Hu, W.-H. Hu, Y. Sun, et al., “Metronomic capecitabine as adjuvant therapy in locoregionally advanced nasopharyngeal carcinoma: a multicentre, open-label, parallel-group, randomised, controlled, phase 3 trial,” The Lancet, vol. 398, no. 10297, pp. 303–313, 2021.

[50] O. Y. Basar, S. Mohammed, M. W. Qoronfleh, and A. Acar, “Optimizing cancer therapy: a review of the multifaceted effects of metronomic chemotherapy,” Frontiers in Cell and Developmental Biology, vol. 12, p. 1369597, 2024.

[51] U. Ledzewicz and H. Schättler, “Application of mathematical models to metronomic chemotherapy: What can be inferred from minimal parameterized models?,” Cancer Letters, vol. 401, pp. 74–80, 2017.

[52] F. Barlesi, L. Deyme, D.-C. Imbs, E. Cousin, M. Barbolosi, S. Bonnet, P. Tomasini, L. Greillier, M. Galloux, A. Testot-Ferry, et al., “Revisiting metronomic vinorelbine with mathematical modelling: A phase i trial in lung cancer,” Cancer Chemotherapy and Pharmacology, vol. 90, no. 2, pp. 149–160, 2022.

[53] M. Strobl, J. Gallaher, M. Robertson-Tessi, J. West, and A. Anderson, “Treatment of evolving cancers will require dynamic decision support,” Annals of Oncology, vol. 34, no. 10, pp. 867–884, 2023.

[54] J. W. T. Yates and H. Mistry, “Clone wars: Quantitatively understanding cancer drug resistance,” JCO Clinical Cancer Informatics, vol. 4, pp. 938–946, 2020. PMID: 33112660.

[55] J. Zhou, C. Liu, Y. Tang, Z. Li, and Y. Cao, “Phenotypic switching as a non-genetic mechanism of resistance predicts antibody therapy regimens,” iScience, vol. 27, Apr 2024.

[56] M. Nagase, S. Aksenov, H. Yan, J. Dunyak, and N. Al-Huniti, “Modeling tumor growth and treatment resistance dynamics characterizes different response to Gefitinib or chemotherapy in non-small cell lung cancer,” CPT Pharmacometrics Syst Pharmacol, vol. 9, pp. 143–152, Mar 2020.

[57] L. Claret, P. Girard, P. M. Hoff, E. Van Cutsem, K. P. Zuideveld, K. Jorga, J. Fagerberg, and R. Bruno, “Model-based prediction of phase iii overall survival in colorectal cancer on the basis of phase ii tumor dynamics,” Journal of Clinical Oncology, vol. 27, no. 25, pp. 4103–4108, 2009. PMID: 19636014.

[58] D. J. Glazar, G. D. Grass, J. A. Arrington, P. A. Forsyth, N. Raghunand, H.-H. M. Yu, S. Sahebjam, and H. Enderling, “Tumor volume dynamics as an early biomarker for patient-specific evolution of resistance and progression in recurrent high-grade glioma,” Journal of Clinical Medicine, vol. 9, no. 7, 2020.

[59] C. Grassberger, I. McClatchy, David, C. Geng, S. C. Kamran, F. Fintelmann, Y. E. Maruvka, Z. Piotrowska, H. Willers, L. V. Sequist, A. N. Hata, and H. Paganetti, “Patient-Specific Tumor Growth Trajectories Determine Persistent and Resistant Cancer Cell Populations during Treatment with Targeted Therapies,” Cancer Research, vol. 79, pp. 3776–3788, 07 2019.

[60] E. Kim, J. S. Brown, Z. Eroglu, and A. R. Anderson, “Adaptive therapy for metastatic melanoma: Predictions from patient calibrated mathematical models,” Cancers, vol. 13, no. 4, 2021.

[61] M. Russo, S. Pompei, A. Sogari, M. Corigliano, G. Crisafulli, A. Puliafito, S. Lamba, J. Erriquez, A. Bertotti, M. Gherardi, F. Di Nicolantonio, A. Bardelli, and M. Cosentino Lagomarsino, “A modified fluctuation-test framework characterizes the population dynamics and mutation rate of colorectal cancer persister cells,” Nature Genetics, vol. 54, pp. 976–984, Jul 2022.

[62] K. Johnson, G. Howard, D. Morgan, E. Brenner, A. Gardner, R. Durrett, W. Mo, A. Al’Khafaji, E. Sontag, A. Jarrett, T. Yankeelov, and A. Brock, “Integrating transcriptomics and bulk time course data into a mathematical framework to describe and predict therapeutic resistance in cancer,” Physical Biology, vol. 18, p. 016001, 2021.

[63] J. A. E. Andersson, J. Gillis, G. Horn, J. B. Rawlings, and M. Diehl, “CasADi – A software framework for nonlinear optimization and optimal control,” Mathematical Programming Computation, vol. 11, no. 1, pp. 1–36, 2019.

[64] A. Wächter and L. T. Biegler, “Line search filter methods for nonlinear programming: Motivation and global convergence,” SIAM Journal on Optimization, vol. 16, no. 1, pp. 1–31, 2005.

[65] H. Hong, A. Ovchinnikov, G. Pogudin, and C. Yap, “Global identifiability of differential models,” Communications on Pure and Applied Mathematics, vol. 73, no. 9, pp. 1831–1879, 2020.

[66] H. Hong, A. Ovchinnikov, G. Pogudin, and C. Yap, “SIAN: software for structural identifiability analysis of ODE models,” Bioinformatics, vol. 35, pp. 2873–2874, 01 2019.

[67] A. F. Villaverde, A. Barreiro, and A. Papachristodoulou, “Structural identifiability of dynamic systems biology models,” PLoS computational biology, vol. 12, no. 10, p. e1005153, 2016.

